# Plectin promotes an aggressive phenotype and represses cytotoxic T cell activity in pancreatic cancer

**DOI:** 10.64898/2026.04.16.718901

**Authors:** Cody L. Wolf, Roxanne K. Ruiz, Sokchea Khou, Robert Cornelison, Edward B. Stelow, Karl M. Kowalewski, Matthew J. Lazzara, Amanda Poissonnier, Lisa M. Coussens, Kimberly A. Kelly

**Author notes:** These authors contributed equally to this work (Cody L. Wolf, Roxanne K. Ruiz, Sokchea Khou, Kimberly A. Kelly). To whom correspondence should be addressed (K.A.K).

## Abstract

**Background:** Pancreatic adenocarcinoma (PDAC) is an abysmal disease, with a poor clinical outcome, largely due to limited life-extending treatments for patients. Notoriously, PDAC displays a T cell-suppressive tumor microenvironment where underlying molecular mechanisms that lead to this phenotype remain poorly understood. To unravel specific mechanisms, we utilized bioinformatic analyses with functional studies and revealed the cytolinker protein plectin (PLEC) as a novel player in regulating the T cell-suppressive tumor microenvironment of PDAC.

**Methods:** Utilizing the TCGA-PAAD dataset, tumor samples were separated by PLEC expression to evaluate patient survival, and pathway analyses associated with increased tumorigenesis. Evaluation of immune infiltration and subsequent immune deconvolution was performed using tidyestimate and CIBERSORTx R packages. Single-cell RNA-seq (scRNA-seq) analysis from 229 PDAC patients was analyzed to investigate signaling dynamics and immune cell infiltration in PLEC^High^ patients. Functional validation was provided using a monoclonal antibody (mAb) against cell surface plectin (CSP) in two murine PDAC models to examine changes in tumor growth and immune cell subset abundance.

**Results:** Our studies revealed that high plectin expression results in an overall worse survival associated with activation of pro-tumorigenic pathways and decreased anti-tumor immune signature in PDAC patients. Analysis via GSEA indicates PLEC^High^ patients display an aggressive phenotype and suppressed pro-inflammatory signaling pathways. Immune ESTIMATE scores were significantly decreased in PLEC^High^ patients, and scRNA-seq analysis revealed that PLEC^High^ tumors display a decrease in anti-tumor CD8^+^ T cells. *In vivo* analyses using an anti-CSP mAb revealed a reduction in tumor growth kinetics compared to IgG control corresponding with a significant increase in proliferating and activated cytotoxic CD8^+^ T cells. Anti-CSP-mediated tumor suppression was inhibited when CD8^+^ T cells were depleted, indicating that anti-CSP treatment is contingent on cytotoxic T cell functionality.

**Conclusion:** Our findings identify plectin as a biomarker of aggressive disease in PDAC, with high plectin expression associated with decreased T cell infiltration, and that treatment with anti-CSP mAb reinstates anti-tumor immunity and decreases tumor volume *in vivo*. These findings position plectin as a high-priority therapeutic target, with the potential to fundamentally reshape immune responses in PDAC and improve outcomes for patients with few remaining options.

## Background

Pancreatic adenocarcinoma (PDAC) is a devastating disease with a median survival of 6 months and a 5-year relative survival rate of only 13% (*1*). In 2026, 67,530 new PDAC cases and 52,740 deaths are estimated in the United States, and it is projected to be the 2^nd^ leading cause of cancer-related deaths by 2030 (*1, 2*). Due to late-stage diagnosis and limited therapeutic options for patients, treatment relies primarily on surgery and chemotherapy (*3–6*). In particular, PDAC has been historically categorized as an “immunologically cold” tumor, making it challenging to treat with immune checkpoint blockade therapy, which has revolutionized cancer treatment across many subtypes (*7, 8*). Understanding the molecular mechanism behind the “immunologically cold” phenotype of PDAC and identifying novel targets that are clinically actionable for most patients is crucial to improving therapeutic outcomes.

Previously, our research group identified plectin as a prognostic biomarker of PDAC (*9*). Plectin is a 500 kDa cytolinker protein within the plakin family encoded by the gene *PLEC* and is known to interact with all three components of the cytoskeleton to stabilize intracellular components, adhesion complexes, and organelles (*10, 11*). In recent years, we and others have reported that plectin is highly expressed in numerous malignancies (*9, 12–22*), including both primary and metastatic cancers (*21*) and is a significant indicator of worse overall survival (*12, 23, 24*). Loss-of-function studies demonstrated that plectin positively regulates classical hallmarks of cancer, including proliferation, migration, adhesion, invasion, and tumor formation (*13, 25–28*). Importantly, our group previously reported that plectin is aberrantly mislocalized to the plasma membrane in aggressive cancers (*9, 21, 25*), and demonstrated that pharmacologic inhibition of membrane-localized plectin (cell surface plectin (CSP)) reduced tumor cell proliferation and migration through MAPK signaling (*22*). While research on plectin’s role in cancer has exponentially grown over the past two decades, the specific mechanism by which plectin promotes cancer progression and decreased survival remains to be fully illuminated.

To further elucidate the role of plectin in PDAC, we performed differential gene expression and pathway analysis using the TCGA-PAAD dataset to identify unique tumorigenic pathways in PLEC^High^ tumors. Here, we uncovered that in addition to roles in proliferation and migration, plectin expression correlates with an “immunologically cold” phenotype in PDAC with suppressed anti-tumoral immune signaling pathways. High plectin expression, as assessed via bulk and scRNA-seq, associated with reduced anti-tumor CD8^+^ T cell infiltration, that is reversed upon pharmacologic inhibition of CSP, indicating that plectin mislocalization to the membrane directly contributes to the immune-cold phenotype in PDAC. Together, these studies identify a novel role for plectin in the tumor microenvironment and represent a new target to unlock the potential of immunotherapies in PDAC.

## Materials and Methods

### Bulk RNA-seq differential gene expression and gene set enrichment analysis

RNA-seq transcriptomic data and clinicopathological features of 185 PAAD patients were acquired from the Genomic Database Commons (https://portal.gdc.cancer.gov/) (*29*) using the Bioconductor package TCGAbiolinks (*30*). Samples were stratified into quartiles based on normalized *PLEC* counts, and differential gene expression was performed using the DESeq2 package (v1.50.2) (*31*). Before running DESeq2, the gene matrix was pre-filtered by removing low counts and for visualization and ranking, log2fold change shrinkage (*32*) and variance-stabilizing transformations were employed (*33–35*). Gene set enrichment analysis (GSEA) was performed using the clusterProfiler (v4.18.4) package (*36, 37*). The following gene set collections were retrieved from the Molecular Signatures Database (MSigDB) via the msigdbr package (v25.1.1) and used for pathway analysis: Hallmark, GO Biological Process, Reactome, KEGG, WikiPathways, and ImmuneSigDB (*38–43*). The following parameters were applied: minimum and maximum gene set sizes of 10 and 1500, respectively; p-value cutoff of 0.05, with p-values adjusted using the Benjamini-Hochberg (BH) procedure. A fixed random seed was set for each collection to ensure reproducibility. Further, we utilized the package pathfindR (v2.7.0) (*44*), an active-subnetwork-oriented pathway enrichment analysis. To reduce redundancy of enriched terms, we implemented hierarchical clustering, which uses a pairwise distance matrix based on the kappa statistics between the enriched terms (as proposed by *Huang et al.* (*45*)) and visualized the results as a bubble plot using the ‘cluster_enriched_terms’ and ‘term_gene_graph’ functions.

### Immune cell analysis and deconvolution

Tumor Purity, Immune, Stromal, and ESTIMATE scores were examined using the tidyestimate package (v.1.1.1) (*46*). For immune deconvolution of bulk RNA-seq samples, Sample-gene matrixes from each quartile were run on the CIBERSORTx webserver (*47*) as mixture files. Relative scores were obtained using the following parameters: LM22 signature gene file, 1000 permutations, and quantile normalization disabled. Results were downloaded using default settings. Bar plots of relative immune fractions and absolute scores were generated using the ggplot2 package (v4.0.2) (*48*). Wilcoxon tests were performed to determine the significance between samples in the first and fourth quartiles. Two-way ANOVA analysis with Tukey test correction was used for multiple comparisons. A confidence interval of 95% was used for all tests.

### scRNA-seq data acquisition and pathways analysis

Single-cell RNA sequencing data were obtained from a previously processed PDAC dataset (https://zenodo.org/records/14199536) (*49*) and analyzed using the Seurat package (v5.4.0) (*50, 51*). Cell populations were subset from the full atlas based on pre-existing cluster annotations into two compartments: malignant ductal cells (DUCTAL, CYCLING DUCTAL) and immune cells (TNK, CYCLING TNK, MYELOID, CYCLING MYELOID, B CELLS, PLASMA, MAST). To enable tractable analysis of the malignant compartment, 30,000 ductal cells were randomly subsampled prior to preprocessing. All cell subsets underwent standard preprocessing including log-normalization, identification of the top 2,000 highly variable features, scaling, and principal component analysis. Dimensionality was assessed via elbow plot and cumulative variance explained, and uniform manifold approximation and projection (UMAP) was computed using the top 30 principal components (*52*).

Patient-level PLEC expression was quantified by computing the mean log-normalized PLEC expression across all ductal cells per patient. PLEC quartile labels were mapped onto immune cells by patient identifier to enable immune compartment comparisons between PLEC^High^ and PLEC^Low^ groups. For formal differential gene expression analysis of the malignant ductal compartment, raw counts were aggregated per patient within each PLEC quartile using Seurat’s AggregateExpression function, with each patient representing an independent biological replicate. This pseudobulk approach was employed to avoid pseudoreplication inherent to cell-level differential expression testing in single-cell data (*53, 54*). Genes with fewer than 10 counts in fewer than 2 samples were excluded prior to testing. Differential expression between quartiles was performed using DESeq2. Mitochondrial and ribosomal protein genes were excluded from analysis. GSEA was performed against the Reactome pathway database using the gseaPathway function from the ReactomePA package (v1.54.0) with a minimum gene set size of 15, a maximum of 500, and a Benjamini-Hochberg adjusted p-value threshold of 0.05. Normalized enrichment scores (NES) were used to assess the direction and magnitude of pathway enrichment, with positive NES indicating enrichment in the higher PLEC quartile relative to Q1. Differences in cell type proportions and module scores between PLEC^High^ and PLEC^Low^ patients were assessed using the Wilcoxon rank-sum test, with multiple testing correction applied using the Benjamini-Hochberg method where applicable.

Immune cell type proportions were computed at the patient level by calculating the percentage of each immune cluster relative to the total immune cell count per patient, with zero counts explicitly assigned to patient-cluster combinations with no observed cells. Total T cell abundance was defined by the proportion of CD3^+^ cells within the TNK and Cycling TNK compartments relative to all immune cells per patient. T cell lineage was assigned at the single-cell level using a multi-gene signature approach to mitigate the effects of CD4 transcript dropout inherent to single-cell RNA sequencing. CD8^+^ identity was scored using mean expression of CD8A, CD8B, GZMB, GZMK, and NKG7, while CD4^+^ identity was scored using mean expression of CD4, IL7R, TCF7, CCR7, FOXP3, and IL2RA, with cells assigned to the lineage with the higher mean signature score. T cell and myeloid subtypes were defined using curated literature-derived gene signatures (*55–57*). Module scores for T cell and myeloid functional states were computed using Seurat’s AddModuleScore function (*58*) and averaged to the patient level for pseudobulk comparisons between PLEC^High^ and PLEC^Low^ groups.

### Subcutaneous Mouse Model

The following procedures were approved and conducted in accordance with the University of Virginia’s Institutional Animal Care and Use Committee (n=4) and at Wuxi Biologics (n=2) for a total of (n=6). KPC915 cells (*59*) were maintained in KPC915 Medium (10% FBS/1% penicillin-streptomycin/DMEM) at 37°C/5% CO^2^ until preparation for injection. Cells were washed and resuspended in DPBS at 2 × 10^7^ cells/mL, then diluted in Matrigel membrane matrix (Fisher, CAT# CB-40234) to 1 × 10^7^ cells/mL The KPC915 tumor cell suspension was subcutaneously injected into one rear flank of female C57BL/6 mice (RRID:IMSR_JAX:000664) in a volume of 100 μL/mouse for a total of 1×10^6^ cells/mouse. Once tumors reached 100 mm^3^, mice were randomized into treatment groups and dosed intravenously in the tail vein with 3.0 mg/kg of αCSP mouse monoclonal antibody (clone 1D7) (*22, 60*) or IgG1κ isotype control (Bio X Cell Cat# BE0083, RRID:AB_1107784), on day 0, 3, and 7. Tumor growth was monitored and measured using calipers. The length (longest diameter) and width (shortest diameter) of tumors were measured approximately twice weekly with calipers, and tumor volume was calculated. Animals were continuously monitored for overall health, body weight, and tumor volume using the standard equation V = (L × W²) / 2. Animals were euthanized if tumor burden was in excess of 1000 mm³, ulcerated, or upon displaying clinical signs of severe illness, including weight loss >10%, inability to get food, extreme lethargy, failure to return to normal activity, or developing clinical signs necessitating euthanasia. Animals were euthanized by CO^2^ followed by cervical dislocation.

### Orthotopic Mouse Model

All animal studies were approved by OHSU’s Institutional Animal Care and Use Committee. Female C57BL/6 mice (RRID:IMSR_JAX:000664) used in this study were purchased from Charles River Laboratories. Mice were anesthetized using continuous isoflurane anesthesia. Abdominal areas were shaved 24 hours before surgery. Buprenorphine SR was administered at 0.01 mg/kg intraperitoneally 15 minutes before surgery started for pain management. The shaved abdominal region of the animals was swabbed with betadine scrub before being cleaned with 70% alcohol. 1.0-1.5 cm left abdominal flank incisions were made, and the spleen and adherent pancreas tissue exteriorized. 4000 tumor cells were implanted into the pancreas in a mixture containing 50% DMEM and 50% Matrigel. PDAC murine KPC4662 (Vonderheide Laboratory) derived from primary tumors of K-rasLsL.G12D/+;Trp53R172H/+;Pdx-1-Cre (KPC) C57BL/6 mice (*61*). 12 to 14 days after implantation, ultrasound was used to detect and measure tumors. Tumors that reached 10-11 mm^2^ were randomized into different groups to initiate treatment. Four doses of αCSP (clone ZB002) mAb or control mIgG2a (Bio X Cell, Cat# BE0085, RRID:AB_1107771) at 10 mg/kg were delivered by IP injection every 72 hours.

For CD8α depletion experiments, tumors that reached 10-11 mm^2^ were randomized into different groups to initiate treatment. 1 mg/kg of anti-CD8α (Bio X Cell Cat# BE0117, RRID:AB_10950145) or rat-IgG2b_k_ (Bio X Cell Cat# BE0090, RRID:AB_1107780) were delivered by IP for the first dose, followed by 500 μg every 5 days for a total of 3 injections. Depletion of CD8^+^ T cells in the peripheral blood was verified via flow cytometry. 72 hours after the first anti-CD8α or rat-IgG2b_k_ injection, mice were treated with 4 doses of 10mg/kg αCSP (clone ZB002) or mIgG2a control every 72 hours, beginning on day four. Tumor size was monitored via ultrasound. 14 days after the start of treatment, animals were euthanized, and tumors were weighed. Measurement of tumor weight and size was unblinded.

### Immune Subset Analysis via Flow Cytometry

Orthotopic tumors were harvested after cardiac perfusion and subjected to enzymatic digestion study endpoints. For subcutaneous tumors, sections of tumor were shipped on ice to Dr. Lisa Coussens at Oregon Health and Science University for immune subset analysis by flow cytometry. Tumors were chopped into small pieces and incubated for 30 minutes with a digestion medium containing 1.0 mg/ml collagenase IV *(Gibco 17104019),* 1.0 mg/ml trypsin inhibitor *(Gibco 17075029),* and 5000U/ml DNaseI *(Roche 10104159001)*. Cell suspensions were washed and incubated with the antibodies listed in Supplementary Table 2. First, cells were incubated with anti-mouse CD16/CD32 *(BDFcBlock 553141)* and live/dead dye *(Molecular Probe Live/Dead Fixable Blue Dead cell stain 50-112-1524)* for 20 minutes. Next, cells were stained with an extracellular antibody cocktail for 30 minutes. For intracellular staining, cells were fixed and permeabilized using the eBioscience Foxp3/Transcription factor staining buffer set (00-5523-00) following manufacturer instructions. Lastly, cells were stained with intracellular antibodies for 30 minutes. Cells were analyzed using Cytek Spectral Aurora. A list of the antibodies, dilutions used, and vendor/catalog information is available in the supplementary material.

### Statistical Analysis

Statistical analysis was performed in R (v4.5.2) (*30*). Results from *in vivo* experiments were analyzed using GraphPad Prism software (version 10, GraphPad Software, Inc., San Diego, CA). Differences between two groups were compared using the Wilcoxon rank-sum test. Multiple medians were compared using the Kruskal-Wallis test, followed by Dunn’s multiple comparisons. For single-cell analyses, patient-level pseudobulk comparisons of immune cell type proportions, T cell and myeloid module scores were assessed using the Wilcoxon rank-sum test. Spearman correlation was used to assess the relationship between PLEC+ ductal cell abundance and immune cell type proportions. Multiple testing correction was applied using the Benjamini-Hochberg false discovery rate method throughout. A two-tailed P value of <0.05 was considered statistically significant for all analyses. The exact value of n within the figures is indicated in figure legends and/or methods sections.

## Results

### High Plectin Expression Is Associated with a Pro-Tumorigenic Phenotype in PDAC

To understand the effects of plectin expression in PDAC, we analyzed 185 patient samples from the TCGA-PAAD dataset (*29*) and stratified patients into quartiles based on their normalized PLEC RNA expression. Patient samples in the first quartile (PLEC^Low^) exhibit the lowest levels of PLEC expression, whereas those in the fourth quartile (PLEC^High^) exhibit the highest levels (Fig. 1A, Supplemental Fig. 1A). Subsequent Cox analysis revealed a significant decrease in survival within PLEC^High^ patients when compared to PLEC^Low^ patients (p = 0.018; Fig. 1B), underscoring the clinical relevance of plectin expression as a prognostic indicator in PDAC.

**Figure 1:**
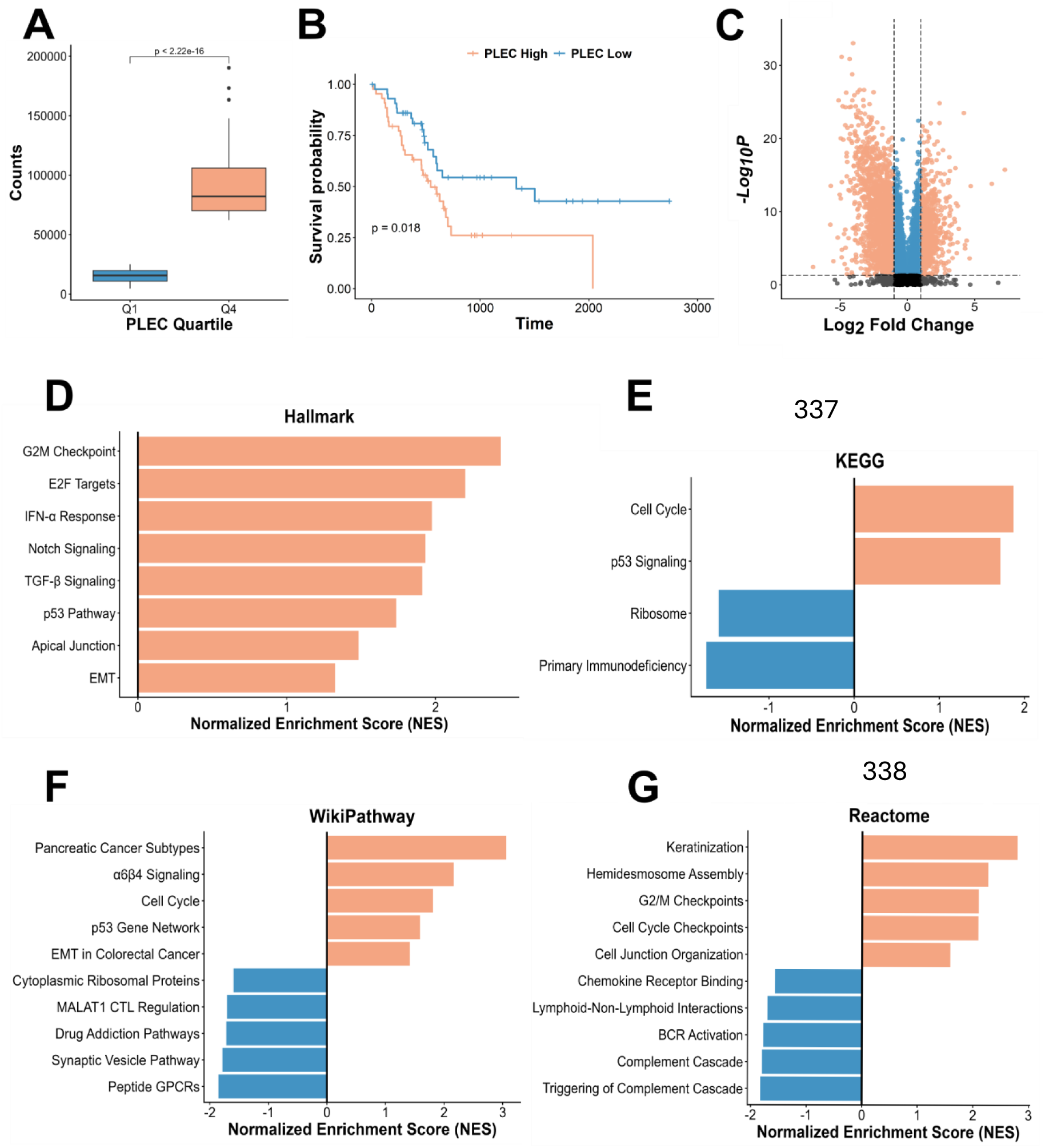
High plectin expression leads to worse survival and an aggressive phenotype in PDAC patients. **(A)** Normalized PLEC RNA expression counts from patients in the TCGA-PAAD dataset, categorized into quartiles. Significance assessed by Kruskal-Wallis test with Dunn’s post-hoc comparisons. **(B)** PLEC^High^ patients (pink; n=44) display significantly worse survival (p=0.018) when compared to PLEC^Low^ patients (blue; n=45). Groups were compared using the log-rank test. **(C)** Volcano plot of differentially expressed genes between PLEC^High^ and PLEC^Low^ samples. Differentially expressed genes were defined by |log₂ fold change| ≥ 2 and adjusted p-value < 0.05. Points are colored as follows: pink, statistically significant, |log₂FC| ≥ 2; blue, statistically significant (p < 0.05) |log₂FC| < 2; grey, |log₂FC| ≥ 2, p-value not significant; black, not significant. GSEA analysis from the Hallmark **(D)**, KEGG **(E)**, WikiPathway **(F)**, and Reactome **(G)**, gene sets from MSigDB identify pathways upregulated (pink) or downregulated (blue) in PLEC^High^ patients compared to PLEC^Low^ patients. PLEC^High^ patients display upregulation in pathways associated with PLEC function, as well as pathways associated with aggressive disease.

After stratifying patient samples, DESeq2 analysis (*31*) was performed to identify differentially expressed genes (DEGs) between the two groups. In total, 1,143 upregulated and 2,823 downregulated DEGs were identified (Fig. 1C, Supplemental Table 1). Gene set enrichment analysis (GSEA) (*38*) across multiple independent gene set databases retrieved from the Molecular Signatures Database (MSigDB) were evaluated: Hallmark (Fig. 1D), KEGG (Fig. 1E), WikiPathways (Fig. 1F), Reactome (Fig. 1G) and GO Biological Processes (Supplemental Fig. 1B) (*38–43*). Consistent with plectin’s established role as a cytolinker, Reactome and GO Biological Processes analyses further confirmed enrichment in structural pathways including Keratinization, Hemidesmosome Assembly, and Cell Junction Organization (Fig. 1G, Supplemental Fig. 1B). Across all four databases, PLEC^High^ samples consistently displayed upregulation of pathways associated with cell cycle progression, including G2M Checkpoint, E2F Targets, p53 Signaling and Cell Cycle Checkpoints (Fig. 1D-G). Upregulation of invasive signaling pathways was also observed, including TGF-β Signaling, Notch Signaling, α6β4 Signaling, Pancreatic Cancer Subtypes, and EMT-associated pathways across multiple databases, collectively reflecting an aggressive and invasive phenotype.

### Plectin Expression Correlates with Decreased Anti-Tumor Immunity in Patients

Surprisingly, when evaluating gene sets, differential plectin expression correlated with immunoregulatory changes, where PLEC^High^ samples associated with suppression of Chemokine Receptor Binding, Immunoregulatory Interactions Between Lymphoid and Non-Lymphoid Cells, and Complement Signaling when compared with PLEC^Low^ samples (Fig. 1F-G). Analysis using the ImmuneSig gene set also identified DEGs associated with increased regulatory T cell (Treg) activity and decreased anti-tumor CD8⁺ T cell responses, indicating that PLEC^High^ patients may exhibit reduced anti-tumor immunity (Fig. 2A). To further evaluate differential gene expression in PLEC^High^ versus PLEC^Low^ patients, we employed pathfindR, an active-subnetwork-oriented pathway enrichment analysis tool that incorporates protein–protein interaction networks in conjunction with gene expression data (*44*). Consistent with GSEA results, pathfindR further illuminated the contribution of PLEC to cancer immunoregulation. PLEC^High^ samples showed downregulation in T cell signaling pathways, including the T cell receptor signaling pathway and Th1/Th2 cell differentiation (Fig. 2B-C). We also observed downregulation of several inflammatory cytokines and their receptors, including IL-2, IL-6, STAT4, and CD3 complex proteins such as CD3ε and CD3δ, which are classically responsible for antigen recognition and T cell activation (*62*) (Fig. 2C). To investigate whether plectin expression directly influenced the composition of the tumor immune microenvironment, we applied the ESTIMATE algorithm via the tidyestimate R package (*46*), which infers tumor purity and the degree of immune and stromal infiltration from bulk gene expression data. PLEC^High^ patients demonstrated significantly increased tumor purity (p = 0.016) and significantly reduced immune score (p = 0.028) and overall ESTIMATE score (p = 0.016) relative to PLEC^Low^ patients (Fig. 2D), while stromal scores showed a decreased trend that did not reach statistical significance in PLEC^High^ patients.

**Figure 2:**
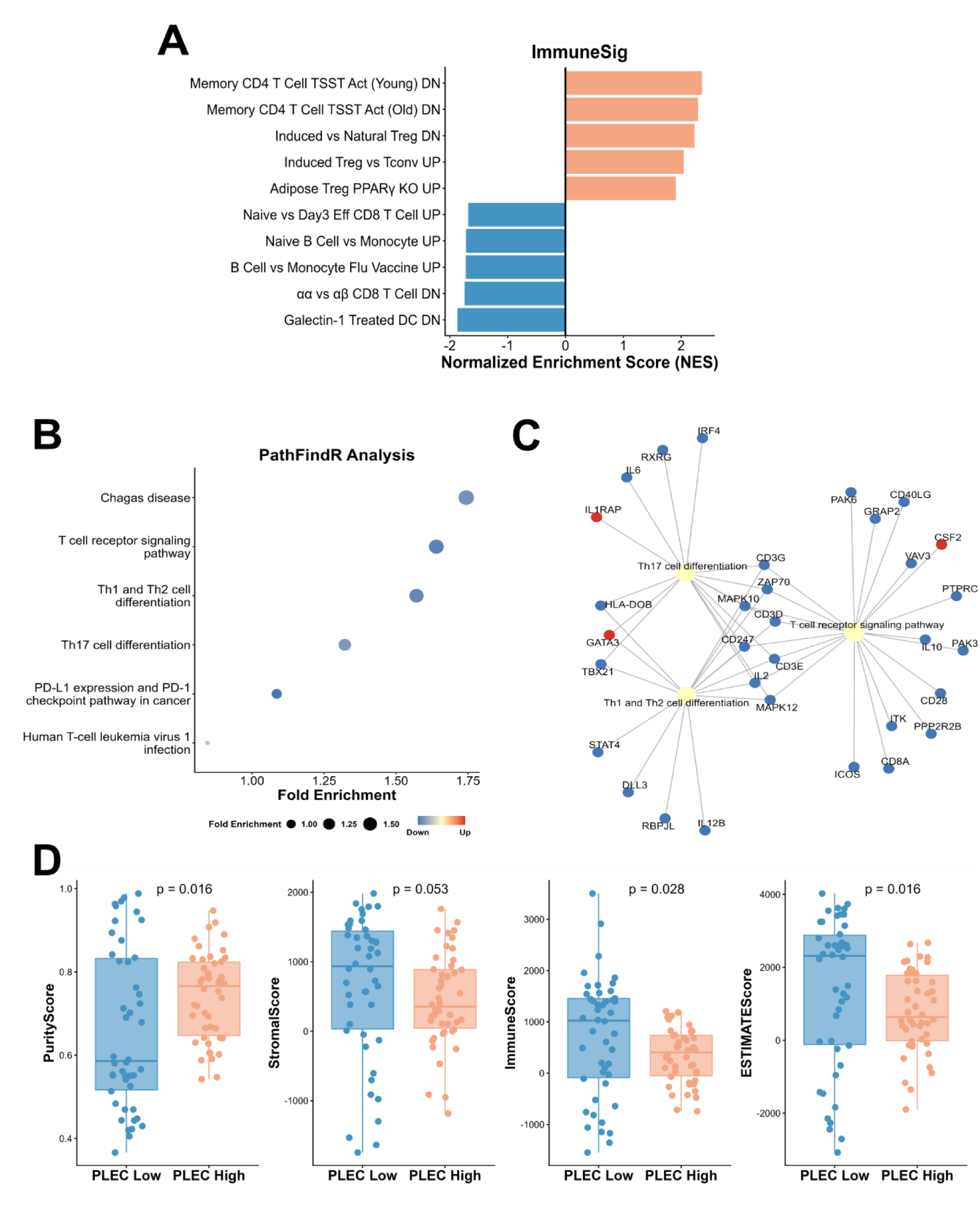
Plectin correlates with an immunosuppressive phenotype in PDAC. **(A)** GSEA analysis from the ImmuneSigDB gene set (MSigDB) suggests PLEC^High^ patients correspond with an immunosuppressive phenotype in cancer. **(B-C)** Pathway analysis using pathfindR after hierarchical clustering identifies suppression of immune-related pathways such as Th1 and Th2 differentiation, and T cell receptor signaling. Notable genes downregulated include CD3 genes, CD28, IL-6, and IL-12. **(D)** ESTIMATE analysis using tidyestimate identifies PLEC^High^ patients display a higher tumor purity score, and a significant decrease in immune and ESTIMATE scores. Wilcoxon rank-sum test.

Based on our bulk transcriptomic analysis, indicating that PLEC^High^ patient samples display decreased anti-tumor immune signaling, we sought to more thoroughly evaluate the immune landscape of PLEC^High^ tumors. To accomplish this within patient tumors, we performed immune deconvolution via CIBERSORTx (*47*) and calculated the relative proportions for 22 immune cell subtypes using the LM22 matrix for each sample across all four quartiles (Fig. 3A, Supplementary Fig. 2A). The proportions of naïve B cells, plasma cells, and CD8⁺ T cells, were lower in PLEC^High^ samples, consistent with an immunosuppressed tumor microenvironment (Fig. 3B). Moreover, PLEC^High^ samples showed a significant increase in M0 macrophages and a slight trending increase in Tregs (Fig. 3C). Taken together, ESTIMATE scores and immune deconvolution analyses indicated that PLEC^High^ tumors correlated with an “immunologically cold” phenotype, while PLEC^Low^ tumors instead correlated with an “immunologically hot” phenotype (*7, 8*). The specific mechanisms by which plectin drives or associated with this altered immune response, however, remain to be elucidated.

**Figure 3:**
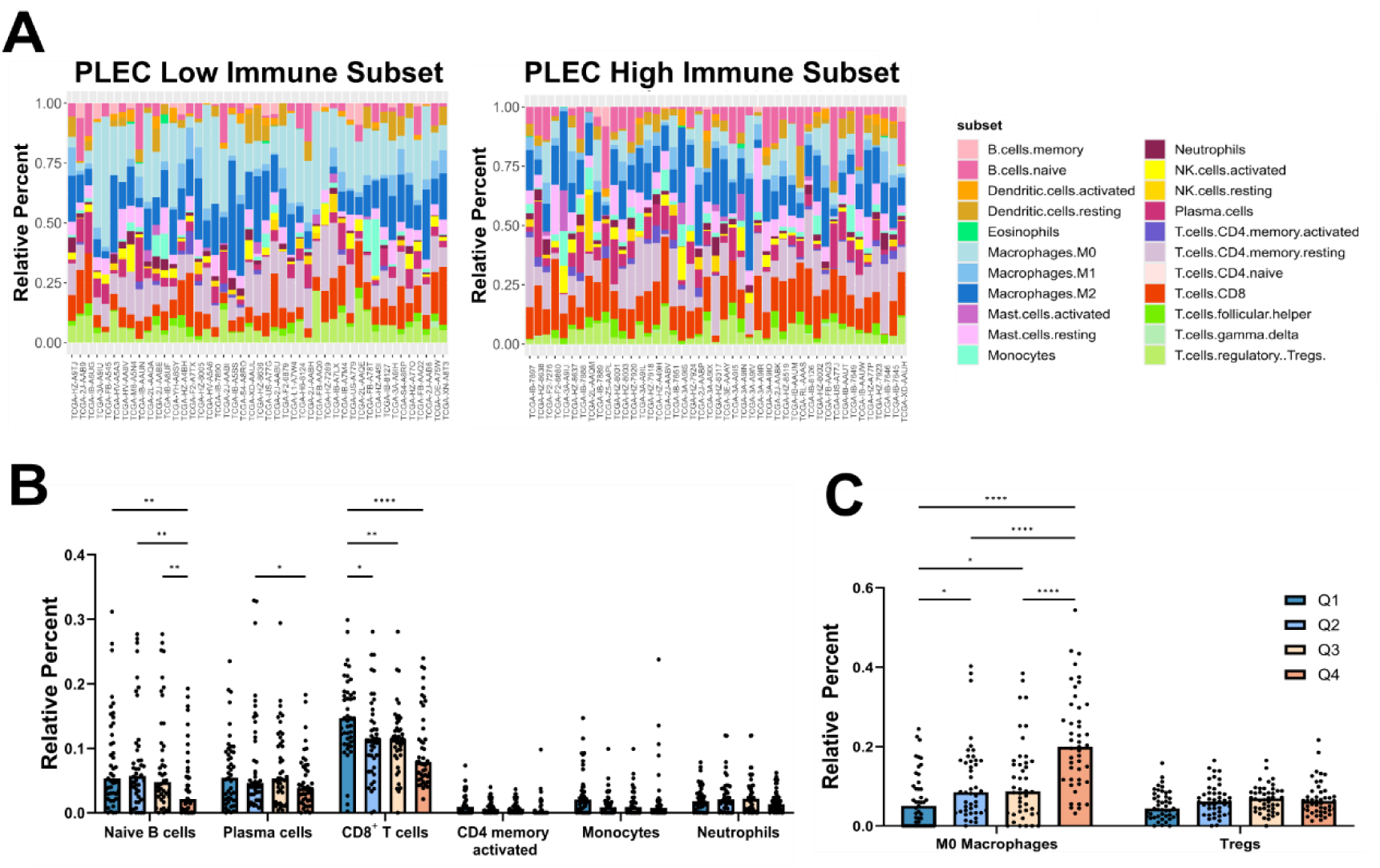
Immune deconvolution reveals that PLEC high patients have a significant decrease in CD8^+^ T cells. PLEC^Low^ and PLEC^High^ patients were evaluated using CIBERSORTx and the LM22 immune signature matrix. (**A**) Bar charts indicate relative percentages of the 22 immune subsets. (**B**) Relative percent scores of the 22 immune subsets for select immune cells reveal a significant decrease in naïve B cells, plasma cells, and CD8^+^ T cells, and an increase in M0 macrophages between quartiles. Kruskall-Wallis test with Dunn’s multiple comparisons (*p<0.05, *p<0.01, ***p<0.001, ****p<0.0001).

### Single-cell Analysis Reveals Plectin Inhibits Cytotoxic T Cell Infiltration

To further evaluate the immunosuppressive nature of plectin in PDAC, we analyzed a previously compiled scRNA-seq dataset containing samples from 229 PDAC patients (*49*). Ductal and Cycling Ductal subsets were isolated and evaluated for PLEC expression (Fig. 4A-B). PLEC expression was abundantly high in both subsets, with a significant increase in the Cycling Ductal group (Fig. 4C), consistent with our previous findings reporting aberrantly high PLEC expression in highly proliferative neoplastic cells (Fig. 1). To evaluate differences in PLEC expression levels, Ductal and Cycling Ductal single cell samples were pseudobulked to the patient level, stratified into quartiles by mean PLEC expression, and GSEA was performed comparing each of the upper three quartiles to the lowest quartile (PLEC^Q1^) samples. DEG’s in each quartile were subsequently evaluated using Reactome pathway gene sets.

**Figure 4:**
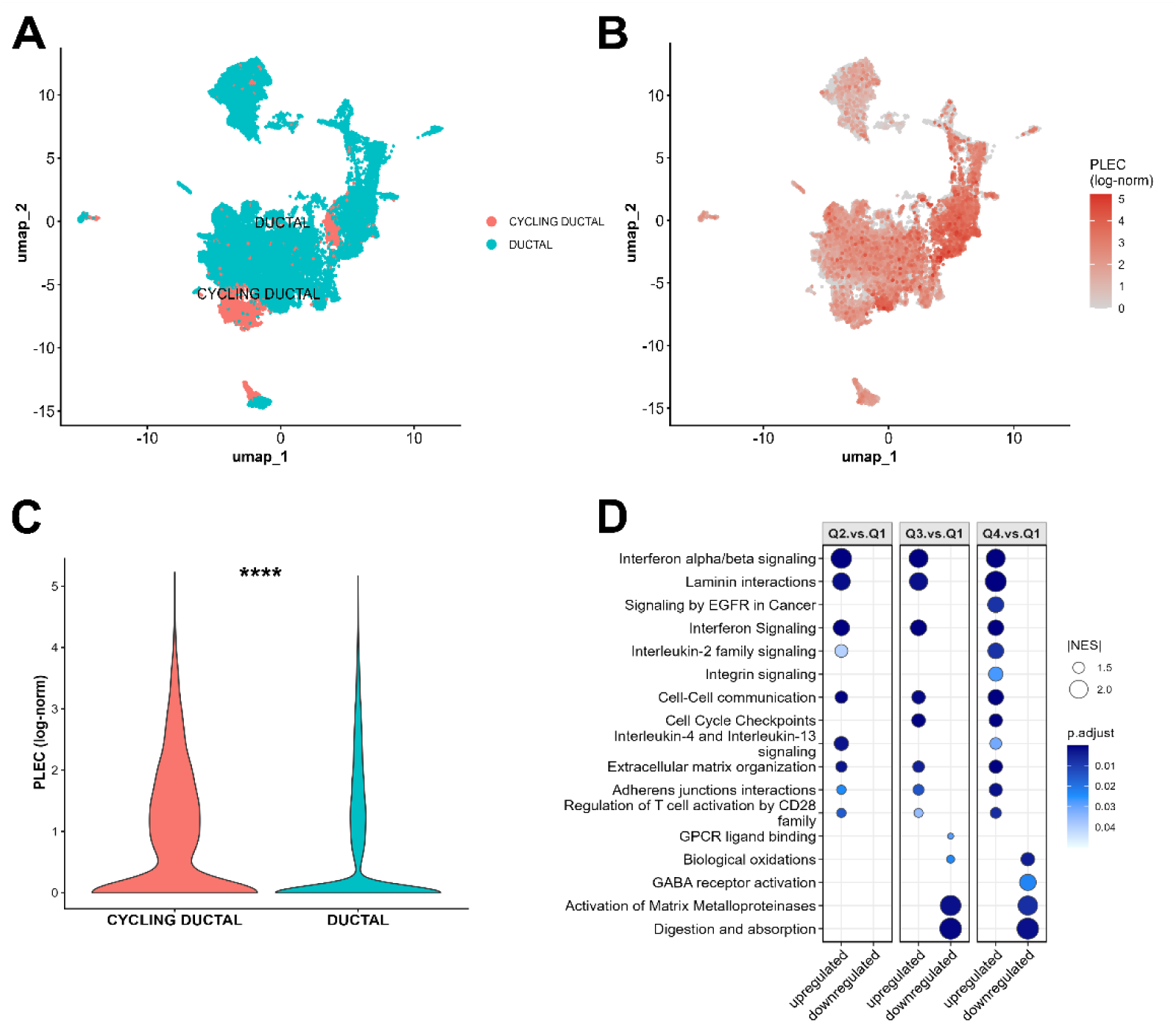
Single-cell analysis correlates plectin expression with highly proliferative and aggressive cells in PDAC subsets. Using a previously annotated dataset (*49*), ductal and cycling ductal cells were evaluated for PLEC RNA expression **(A-B)**. **(C)** PLEC expression was significantly upregulated in cycling ductal cells compared to ductal cells, suggesting high PLEC expression correlates with a more aggressive phenotype. Wilcoxon rank-sum test. **** p<0.0001. **(D)** GSEA analysis was performed against the Reactome database (MSigDB) to identify differentially enriched gene set pathways. Pseudobulked single-cell RNA-seq counts were quartiled by PLEC expression. Dot size reflects normalized enrichment score (NES), and color corresponds to BH-adjusted p-value. Across all quartiles, gene sets corresponding to an aggressive tumor phenotype were observed, as well as pathways suggesting immune suppression.

As in our bulk analysis, high PLEC expression correlated with upregulated genes related to previously known plectin function, such as Laminin Interactions, Integrin Signaling, Cell-Junction Organization, and Cell-Cell Communication pathways (Fig. 4D). High plectin was also found to correlate with pro-tumorigenic pathways, including Cell Cycle Progression and Extracellular Matrix Organization across all quartiles, with effects more pronounced in PLEC^Q4^ cells. Excitingly, both Interferon Signaling and Regulation of T cell Activation were upregulated across all quartiles (Fig. 4D). Notably, the concurrent upregulation of Interferon Signaling and Regulation of T cell Activation pathways in plectin high tumor cells indicated a paradoxical immune activation state, wherein tumor-intrinsic interferon signaling may drive expression of immune checkpoint ligands such as PD-L1, ultimately dampening rather than promoting cytotoxic T cell function (*63, 64*).

We further explored the relationship between plectin and tumor-immune cell communication by analyzing the single-cell immune compartment. Annotated immune cells were normalized for compartmentalization via UMAP dimensionality reduction (Fig. 5A), and expression of canonical markers for distinct immune cell populations in the TNK and Myeloid compartments, including T cell markers (CD3ε,CD3δ), myelomonocytic markers (CD14, LYZ) and natural killer (NK) cell markers (NCAM1, GNLY; Fig 5B) (*55–57*). Comparison of mean immune cell proportions between PLEC^Low^ and PLEC^High^ revealed a marked decrease in B cells and TNK Cells, alongside a concurrent increase in Cycling Myeloid Cells, Myeloid Cells, and Mast Cell compartments in PLEC^High^ patients (Fig. 5C). Given the observed shifts in broad immune compartments, we next interrogated the TNK and myeloid populations in greater detail. Stratification of TNK and cycling TNK cells by total T cell population and by CD4 and CD8 expression revealed a trending reduction in T cells (Fig. 5D), significant increase in CD4⁺ cells (p < 0.001) and a significant decrease in CD8⁺ cells (p < 0.01) within the PLEC^High^ cohort (Fig. 5E), while no significant change was observed when stratifying myeloid cell subtypes (Supplemental Fig. 3A). Finally, assessment of T cell phenotypic markers across PLEC^High^ and PLEC^Low^ patients revealed a modest, non-significant increase in both exhausted T cells (T_ex_) and Tregs in PLEC^High^ patients (Supplemental Fig. 4), further supporting the notion that elevated plectin expression promotes a T cell-suppressive tumor microenvironment characterized by reduced cytotoxic cell recruitment.

**Figure 5:**
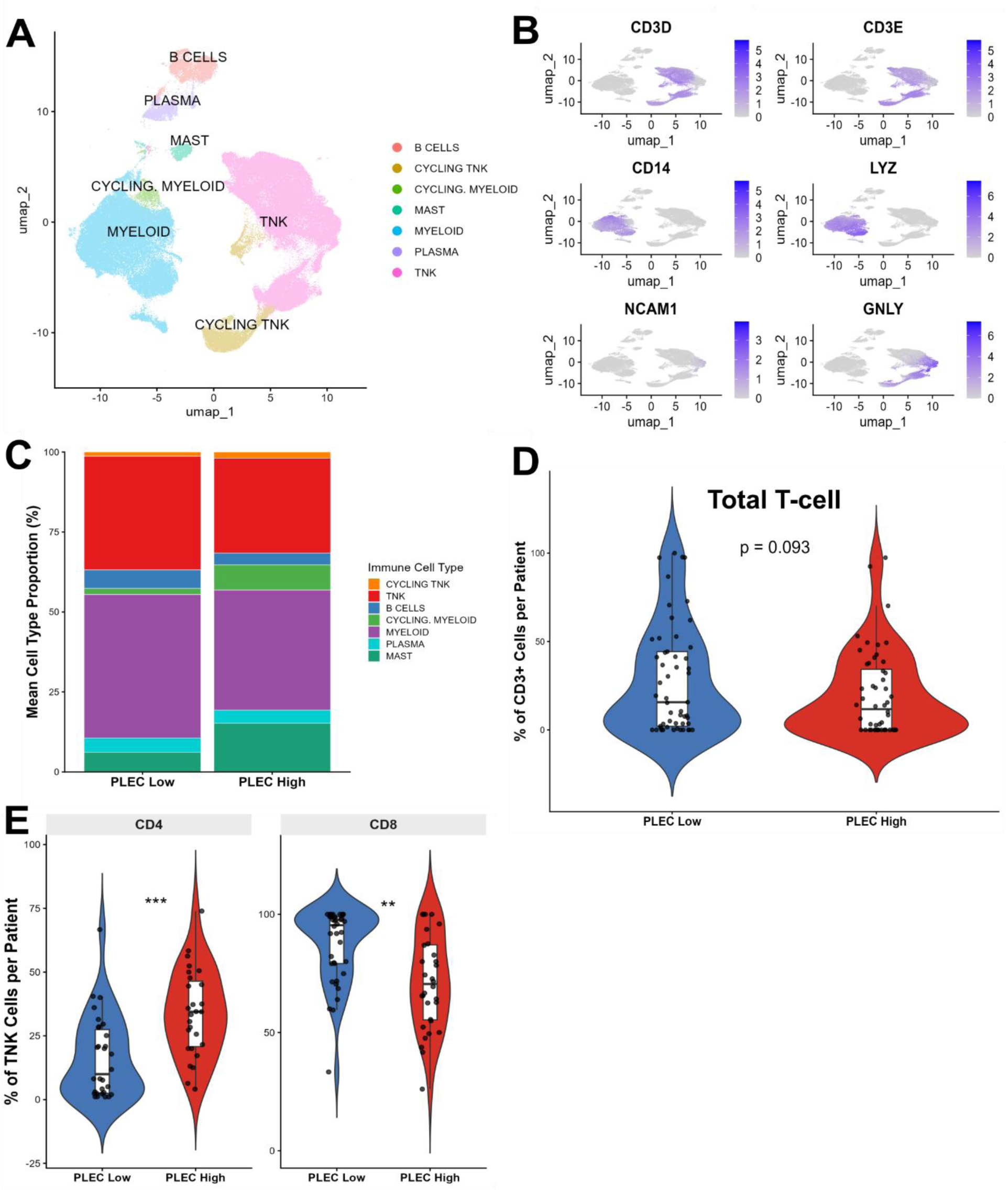
Immune subset analysis identifies plectin high patients displaying reduced CD8^+^ T cells in PDAC patients. **(A)** UMAP plot showcasing immune cell signatures within the immune compartment of the *Loveless et al* dataset (*49*). **(B)** Expression of specific gene signatures for distinct immune cells were evaluated by feature plot within the immune subset. T cells were identified by CD3ε and CD3δ expression, myeloid cells by CD14 and LYZ and NK cells by NCAM1 and GNLY markers. **(C)** Mean cell type proportion of each immune subset was evaluated between PLEC^Low^ and PLEC^High^ patients, revealing a decrease in the myeloid and TNK compartments. **(D)** Patient-level pseudobulk quantification of total T cell abundance via RNA expression of CD3 in TNK and Cycling TNK clusters, reveals a trending decrease in T cell population in PLEC^High^ patients when compared to PLEC^Low^ patients. **(E)** Within the TNK and Cycling TNK compartments, PLEC^High^ patients display an increase in the proportion of CD4^+^ T cells and a corresponding decrease in CD8^+^ T cells relative to PLEC^Low^ patients, indicating a shift in the cytotoxic to helper T cell balance. Wilcoxon rank-sum tests ** p<0.01, *** p <0.001

### Targeting Cell Surface Plectin Results in Increased CD8^+^ T cell infiltration *in vivo*

To experimentally examine whether plectin contributes to establishment of a T cell-excluded immune microenvironment, and if an anti-tumorigenic immune response could be reinstated by therapeutically targeting plectin, we sought to leverage a previously evaluated neutralizing monoclonal antibody specific to Cell Surface Plectin (CSP) (*22*). To facilitate the accuracy of longitudinal tumor measurements, we utilized two different murine models of pancreatic cancer. KPC915 cells were subcutaneously injected into the flank of C57/BL6 mice. Once tumors reached 100 mm^3^ in tumor volume, mice were randomized into two treatment groups: 1) αCSP mAb at 9.0 mg/kg/wk, and 2) mIgG isotype control mAb at 9.0 mg/kg/wk. Mice received a total of three doses in the first week, were monitored for tumor growth kinetics, and tumors harvested for immune subset analysis. αCSP treatment resulted in a >60% tumor regression at day 10, whereas treatment with an IgG control antibody did not alter tumor growth (Fig.6A). To examine changes in immune subsets after treatment, tumors were digested to generate single-cell suspensions and analyzed via flow cytometry using validated lineage-selective antibodies (Supplementary Table 2). Global immune infiltrates were evaluated as a percentage of live CD45^+^ cells. As predicted from our *in-silico* analysis, we found a significant increase in cytotoxic CD8^+^ T cells in tumors from αCSP-treated animals when compared to IgG-treated controls (Fig. 6B).

**Figure 6:**
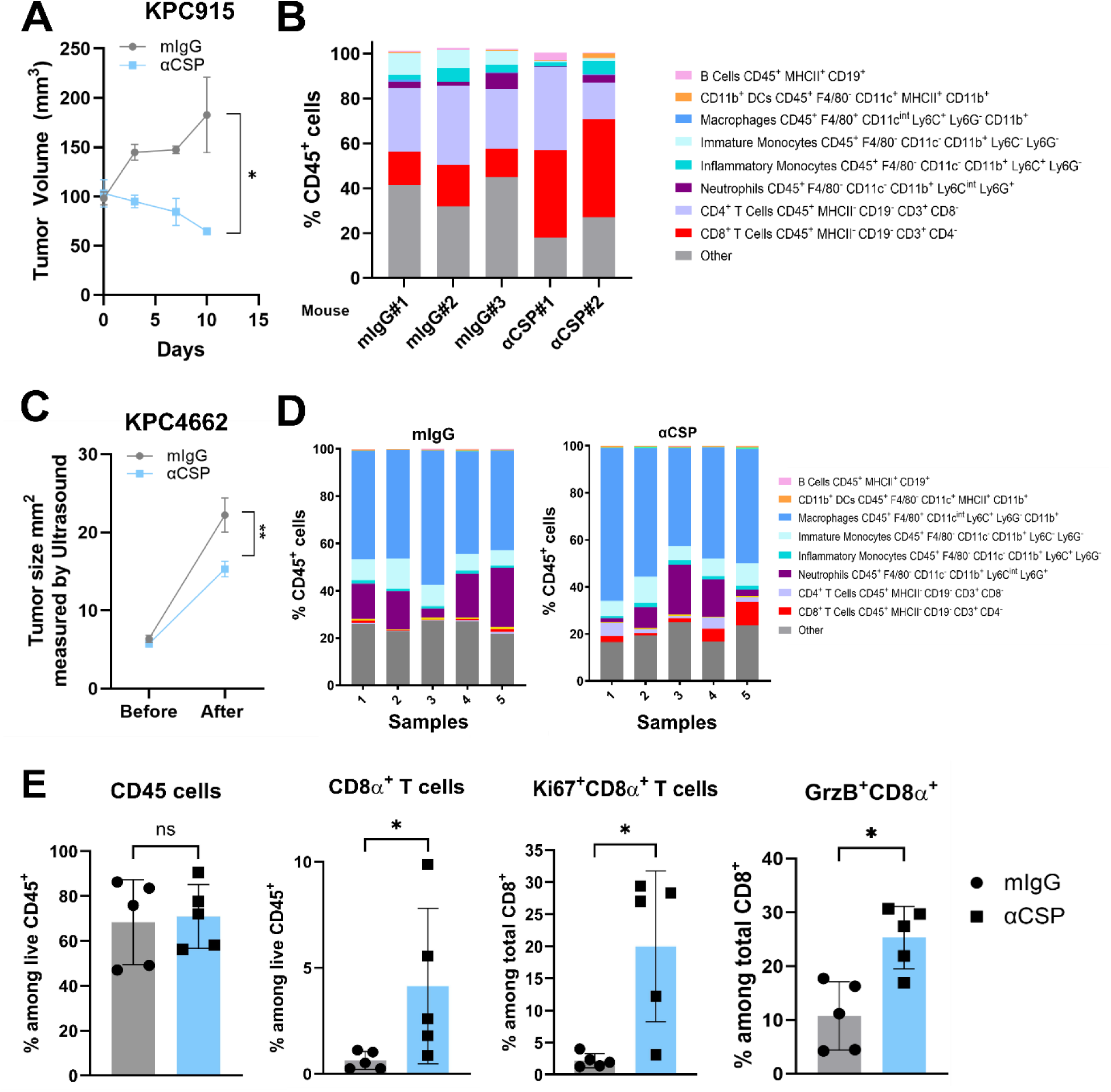
CSP inhibition causes tumor regression and CD8^+^ T cell infiltration. **(A)** Tumor growth analysis from mice injected s.c. with KPC915 and subsequently treated with mIgG isotype control or 9.0 mg/kg/wk αCSP show a significant decrease in tumor volume over 10 days. **(B)** Tumors treated after one (αCSP#1) or two doses (αCSP#2) of αCSP or one dose of mIgG isotype control (mIgG#1, 2,3) were digested to generate a single-cell suspension and stained with multiple antibodies via flow cytometry. Global immune infiltrates shown as a percentage of CD45^+^ cells. Each bar represents an individual tumor. **(C)** Orthotopic xenograft tumor cell transplantation of KPC4662 cells into the head of the pancreas and subsequent treatment of mIgG or αCSP (10 mg/kg) show significant decrease in tumor size as measured by ultrasound. **(D-E)** After single-cell digestion, mice treated with αCSP show a significant increase in CD8^+^ T cells, including Ki67^+^ and Granzyme B^+^ (GrzB^+^) CD8^+^ T cells, showcasing proliferative anti-tumor activity after αCSP treatment. Wilcoxon rank-sum test. *p<0.05.

To investigate the actions of plectin inhibition in a more clinically relevant model, we used a second murine-derived PDAC cell line, KPC4662, and implanted tumors orthotopically in the pancreas of C57BL/6 mice. Tumor growth was monitored by ultrasound (Supplementary Fig. 5A), and once tumors reached 10 mm^2,^ mice were randomized into two treatment groups: 1) αCSP mAb at 10 mg/kg and 2) IgG mAb isotype control at 10 mg/kg. Treatment occurred every three days, and 48 hours after the last treatment on day 10, tumors were harvested and measured with calipers (Fig. 6C). Treatment with αCSP at 10 mg/kg resulted in a significant decrease in tumor weight (p < 0.01) at 12 days post start of treatment. To examine changes in immune subsets after treatment, tumors were digested to generate single-cell suspensions (Fig. 6D). As predicted from our CIBERSORTx analysis, scRNA-seq analysis, and our subcutaneous model, αCSP treatment led to a trending increase in CD3^+^ T cells (p-value = 0.0556) and significantly increased the presence of CD4^+^ and CD8^+^ T cell populations (p < 0.05) (Fig 6D, Supplemental Fig. 5B). Further analysis revealed that the CD4^+^ and CD8^+^ T cell populations in the PLEC-inhibited tumors were proliferating and actively cytotoxic, as evidenced by expression of Ki-67 (p< 0.05) and Granzyme B (p < 0.05) (Fig. 6E, Supplemental Fig. 5B) (*65*).

### Tumor regression from plectin inhibition is CD8^+^ T cell-dependent

Since targeting plectin with an αCSP mAb led to an increased presence of CD8^+^ T cells and subsequent tumor suppression, we sought to determine whether the anti-tumoral effects were CD8^+^ T cell-dependent. Mice with orthotopically implanted KPC4662 tumors were randomized into four treatment groups: 1) mIgG + rat IgG, 2) mIgG + αCD8α, 3) rat IgG + αCSP, and 4) αCD8α + αCSP. Treatment was initiated once tumors reached 10 mm^2^, and CD8^+^ T cell depletion occurred two days before administering αCSP or mIgG isotype control. 48 hours after the last treatment, tumor weight and volume were measured (Fig. 7A). As expected, depletion of T cells alone (mIgG + αCD8α) or both isotype control antibodies (mIgG + rat IgG) did not influence tumor weight or volume. Rat IgG+αCSP treatment significantly reduced tumor weight compared to mIgG + rat IgG or mIgG + αCD8α treatment groups (p < 0.05). Importantly, CD8^+^ T cell depletion before αCSP (αCD8α + αCSP) abrogated the anti-tumoral effects of αCSP (Fig. 7B-C). Cytometric analysis after pre-gating on CD3ε^+^ cells (Fig. 7D) as well as measuring the fraction of CD8^+^ T cells among all CD45^+^ cells (Fig. 7E), established that CD8^+^ T cells were, in fact, depleted by the treatment endpoint. Further, the fraction of CD8^+^ T cells among all CD45^+^ peripheral blood mononuclear cells (PBMCs) before and after treatment confirmed depletion efficiency (Fig. 7F). These results demonstrate that the anti-tumoral effects of αCSP treatment are dependent on CD8^+^ T cell cytotoxicity and affirm a critical role for CSP impacting tumor immune evasion.

**Figure 7:**
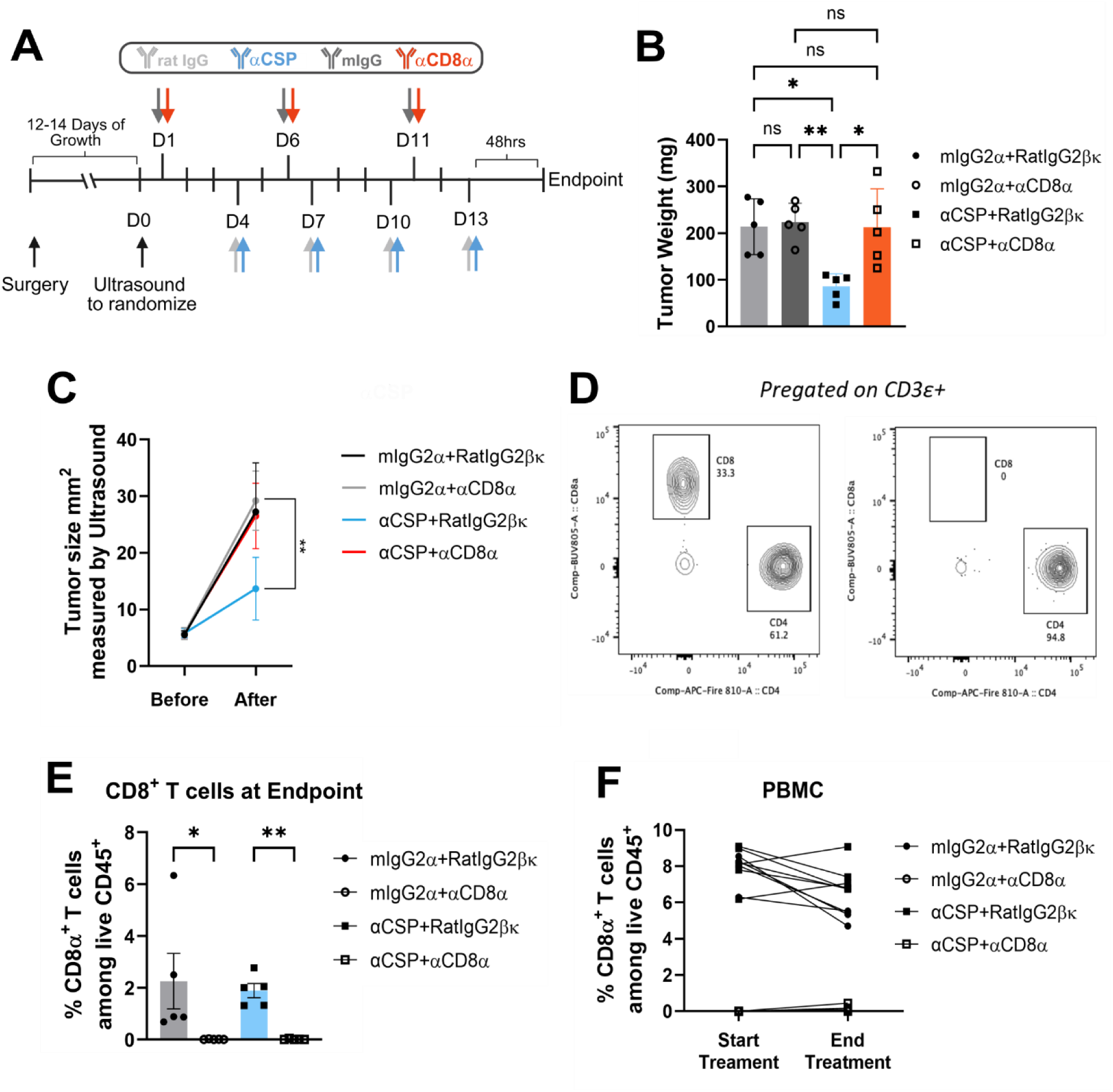
Anti-CSP-mediated tumor regression is CD8^+^ T cell-dependent. (**A**) Treatment schema. Mice were orthotopically implanted with murine KPC4662 cancer cells and subsequently treated with either rat IgG, mouse IgG, αCSP or αCD8 on days specified (n=5 for all groups). (**B**) Tumor weight at study endpoint shows significant decrease in αCSP treated mice and recovery of αCSP and αCD8 treated mice. (**C**) Tumor size measured by ultrasound confirms a significant decrease in tumor growth in αCSP treated mice. (**D**) Flow cytometry analysis of CD8^+^ T cells, pre-gated on CD3ε^+^ T cells, before treatment and after treatment show a complete lack of CD8^+^ cells in mice treated with αCD8. (**E**) CD8^+^ T cell population as a percentage of CD45^+^ cells per treatment group. (**F**) Individual measurements of CD8^+^ T cells as a percentage among live CD45^+^ cells in PBMC before and after treatment. Error bars represent standard deviation. Kruskal-Wallis test. *p<0.05, ** p <0.01.

## Discussion

While personalized medicine has revolutionized cancer treatment and dramatically improved survival across many malignancies, the identification of clinically actionable targets that improve PDAC survival remains an unmet clinical need. This year, for the first time, the overall 5-year survival rate for all cancer patients has reached 75%; yet for PDAC patients, this figure remains a disheartening 13%, a disparity that underscores the urgent need for new therapeutic strategies (*1, 4, 6*). This poor prognosis is driven in large part by the paucity of effective systemic therapies, leaving surgery and chemotherapy as the primary treatment options for most patients. Compounding this challenge, PDAC has long been characterized as an immunologically "cold" tumor, rendering it largely refractory to immune checkpoint blockade therapies that have transformed the treatment landscape for other malignancies (*6, 7*). Identifying mechanisms that underlie this immunosuppressive phenotype, and therapeutic strategies capable of converting cold tumors to immunologically active ones, represents a critical frontier in PDAC research. Here, we identify plectin as a novel immune-modulatory driver of the immunosuppressive tumor microenvironment in PDAC.

Our group previously employed a functional proteomic approach and revelaed that plectin is aberrantly mislocalized to the cell surface of PDAC and other cancers, while remaining cytoplasmic in healthy tissue (*9, 20, 21*). Plectin expression has since been associated with worse overall survival across numerous malignancies, implicating it as an important mediator of tumorigenesis (*13, 14, 16, 18, 23, 24, 27*). Building on these findings, we now demonstrate that elevated plectin expression is directly associated with poor patient survival and an aggressive transcriptomic phenotype in the TCGA-PAAD dataset (Fig. 1). Critically, PLEC^High^ patients displayed an “immune-cold” gene expression signature (Fig. 2), correlating with significantly decreased immune infiltration scores and a marked reduction in B cells and CD8⁺ T cells upon immune deconvolution (Fig. 3).

To extend these findings to a higher-resolution platform, we leveraged a combined single-cell RNA-seq dataset previously annotated by Loveless et al (*49*). PLEC expression was most strongly enriched in Cycling Ductal cells, a transcriptionally distinct ductal subset characterized by high expression of STMN1 and HMGB2, markers associated with aggressive tumor biology (Fig. 4A-C). GSEA of ductal cell populations further confirmed an aggressive, pro-tumorigenic transcriptional program in PLEC^High^ ductal cells, consistent with our bulk transcriptomic analysis, including upregulation of Laminin Interactions, Integrin Signaling, Cell Cycle Progression, and Extracellular Matrix Organization pathways (Figs. 1D-G, 4D). Notably, Interferon Signaling and Regulation of T cell Activation were also upregulated in PLEC^High^ ductal cells across all quartiles, presenting an apparent paradox given the immune-excluded phenotype observed in our single-cell immune analysis.

Interrogation of the immune compartment revealed that PLEC^High^ patients displayed a significant reduction in total T cell infiltration and a concurrent expansion of myeloid populations (Fig. 5). Further stratification by canonical immune markers demonstrated a significant increase in CD4^+^ T-helper cells (p < 0.001) and a significant decrease in CD8^+^ cytotoxic T cells (p < 0.01), corroborating the immunosuppressive phenotype observed in bulk analyses (Fig. 5E). T cells present in PLEC^High^ tumors displayed slightly elevated exhaustion module scores, marked by upregulation of checkpoint inhibitors, indicating that residual T cells are functionally impaired rather than immunologically active (Supplemental Fig. 4).

Rather than reflecting productive anti-tumor immunity, the upregulation of tumor-intrinsic Interferon Signaling in PLEC-high ductal cells likely represents a paradoxical immune evasion mechanism. Interferon signaling is well established to play pleiotropic roles in cancer, while acute interferon activation promotes anti-tumor immunity, chronic or tumor-intrinsic interferon pathway activation can drive transcriptional resistance to immune checkpoint blockade through sustained upregulation of PD-L1, IDO1, and other immunosuppressive mediators (*63, 64*). Prolonged interferon exposure further promotes T cell exhaustion through upregulation of inhibitory receptors (*66, 67*), consistent with the exhaustion phenotype we observe in PLEC^High^ TNK cells.

Given that plectin has been found on the cell surface of cancer cells (CSP) and that CSP has been identified as a viable therapeutic target in various cancers (*9, 20, 22*), we hypothesized that plectin-mediated immune modulation was a direct consequence of CSP expression. Inhibition of CSP using a previously generated monoclonal antibody resulted in significant tumor volume reduction in two independent murine PDAC models (Fig. 6). Immune profiling of treated tumors revealed a significant increase in CD8⁺ T cells relative to IgG controls (Fig. 6B,D), accompanied by robust expression of Ki-67 and Granzyme B, indicating that these cells were cytotoxically active (Fig. 6E). This finding is particularly striking given that durable induction of T cell–mediated tumor killing remains one of the most significant barriers to effective PDAC treatment (*8, 68*).

To formally establish whether these anti-tumoral effects were CD8⁺ T cell dependent, we employed an *in vivo* depletion strategy in orthotopically implanted KPC4662 tumor-bearing mice (Fig.7). CD8⁺ T cell depletion prior to αCSP administration completely abrogated its anti-tumoral effects, while depletion alone had no independent effect on tumor growth (Fig. 7B,C). Depletion efficiency was confirmed both intratumorally and in PBMCs (Fig. 7D,F), ensuring that the loss of therapeutic efficacy was attributable to CD8⁺ T cell absence rather than incomplete depletion. Collectively, these data demonstrate that αCSP anti-tumoral activity is CD8⁺ T cell-dependent and position CSP as a functionally significant mediator of immune evasion in PDAC, one whose pharmacologic inhibition is sufficient to restore cytotoxic T cell killing in a disease where T cell exclusion remains a defining barrier to effective immunotherapy

Taken together, our findings establish plectin as a novel biomarker of immune exclusion and a compelling therapeutic target in PDAC. Although our findings are the first to definitively establish plectin as a negative immunomodulator in PDAC, independent corroboration has recently emerged. *Ge et al*. employed a machine learning-based approach to identify biomarkers distinguishing immunologically “cold” and “hot” PDAC tumors and independently identified plectin as a negative immune regulator (*69*), providing orthogonal validation of our central hypothesis. An important open question is whether the pro-tumorigenic and immunosuppressive effects observed here are driven by global plectin overexpression, aberrant CSP mislocalization, or both. Disentangling these contributions will be critical to understanding the precise mechanisms by which plectin shapes the tumor immune microenvironment and to developing the most effective therapeutic approach. Further analysis of patient tumor samples for specific immune cell subtypes in PLEC^High^ samples will help unravel the precise mechanisms by which CSP represses CD8^+^ T cell infiltration. Elucidating the precise mechanisms of CSP-mediated mediates immune suppression will deepen our understanding of tumor microenvironment dynamics and may uncover additional therapeutic vulnerabilities in this devastating disease. In particular, directly examining regulatory T cell and exhausted T cell populations via flow cytometry or spatial transcriptomics will be critical to fully characterizing CSP-mediated immune modulation and its therapeutic implications in PDAC.

## Conclusions

In summary, this study identifies plectin as a novel negative immunomodulator in PDAC, demonstrating that elevated plectin expression correlates with a T cell-suppressive tumor microenvironment characterized by reduced CD8⁺ cytotoxic T cell infiltration, and broad suppression of pro-inflammatory immune signaling. Pharmacologic targeting of cell surface plectin via αCSP monoclonal antibody was sufficient to restore CD8⁺ T cell-mediated anti-tumor immunity and significantly reduce tumor burden *in vivo*, in a manner that was strictly CD8⁺ T cell dependent. These findings are of particular clinical relevance given the notorious resistance of PDAC to existing immunotherapies and the continued lack of effective treatment options for patients with advanced disease. By establishing CSP as both a biomarker of immune exclusion and a functionally actionable therapeutic target, this work provides a compelling rationale for the continued development of anti-CSP therapies, either as standalone treatments or in combination with existing immune checkpoint inhibitors, with the potential to fundamentally shift the treatment landscape for one of the most lethal malignancies.

## Supplemental Material

**Supplemental Figure 1:**
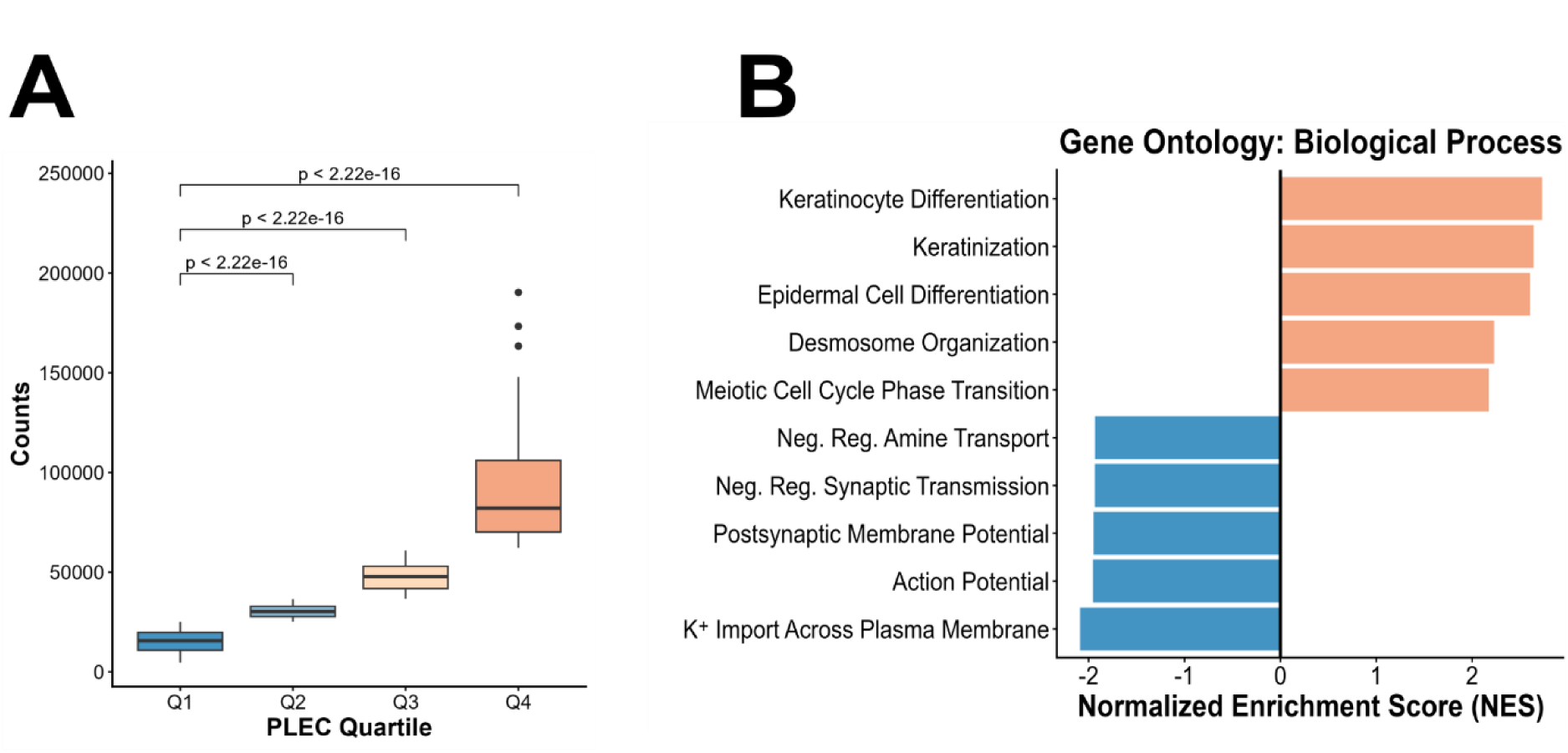
Plectin expression corresponds with a more aggressive phenotype in PDAC. **(A)** Patients within the TCGA-PAAD dataset were categorized into quartiles based on normalized *PLEC* counts. Kruskal-Wallis test **(B)** GSEA of Gene Ontology: Biological Processes of upregulated (pink) and downregulated (blue) gene sets.

**Supplemental Figure 2:**
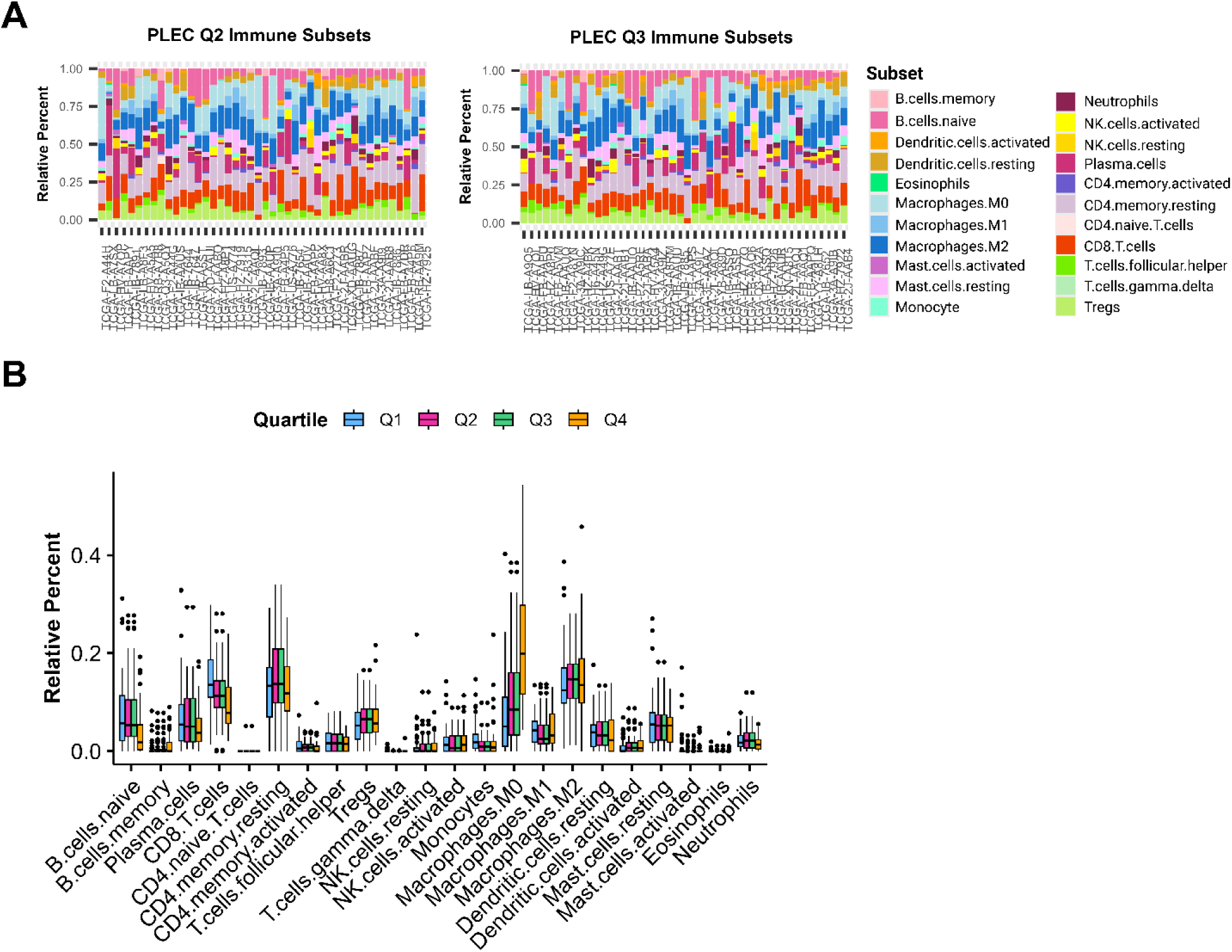
Immune deconvolution of bulk PDAC data. PDAC patients from the second and third quartiles were run on the CIBERSORTx webserver using the LM22 immune signature matrix. (**A**) Bar charts indicate the relative percentages of the 22 immune subsets from Q2 and Q3 expressing samples. (**B**) Relative percent all 22 immune subsets from each quartile.

**Supplemental Figure 3:**
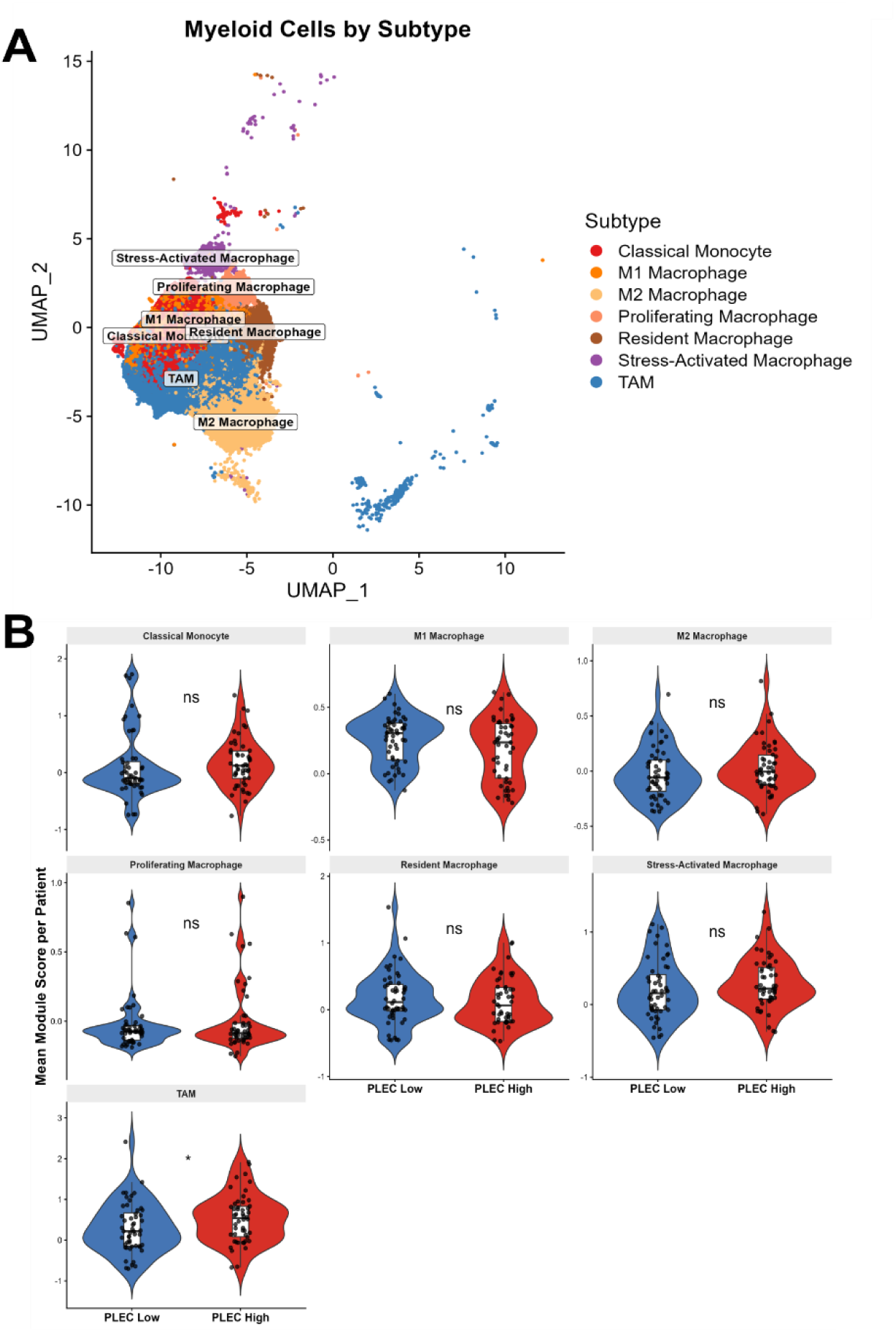
Myeloid subtyping in PLEC^High^ vs PLEC^Low^ samples. **(A)** UMAP plot of myeloid subtypes based on genetic signatures. **(B)** Module scores of each myeloid subtype stratified by PLEC expression. Wilcoxon rank-sum test. ns = not significant.

**Supplemental Figure 4:**
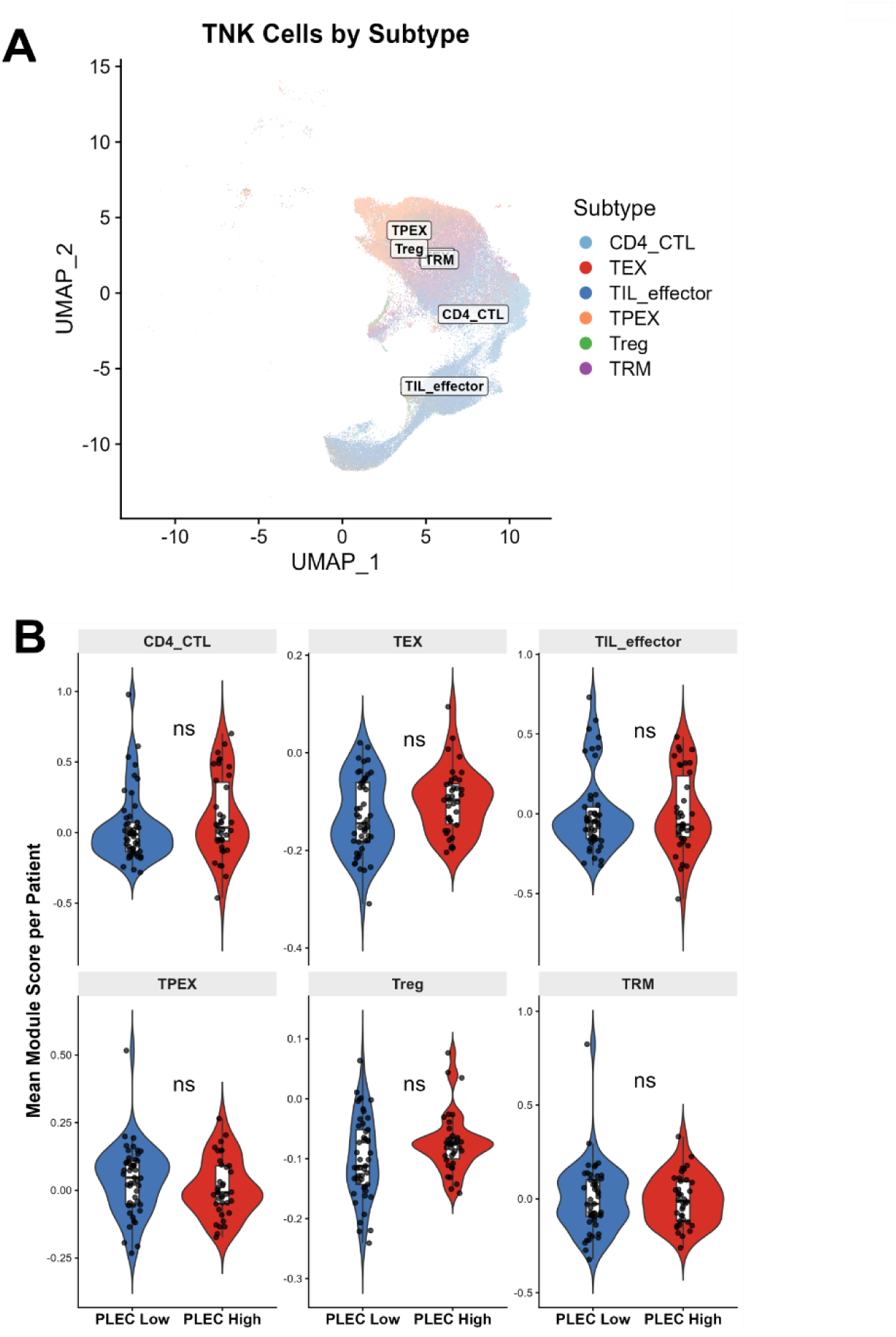
T-cell subtyping in PLEC^High^ vs PLEC^Low^ samples. **(A)** UMAP of distinct T cell populations based on genetic signatures: terminally exhausted T cells (TEX), progenitor exhausted T cells (TPEX), effector TILs (TIL effector), cytotoxic CD4^+^ T cells (CD4 CTL), regulatory T cells (Treg), and tissue-resident memory T cells (TRM)**. (B)** Module scores of each T cell subtype in PLEC^Low^ and PLEC^High^ patients. Wilcoxon rank-sum test. ns = not significant.

**Supplemental Figure 5:**
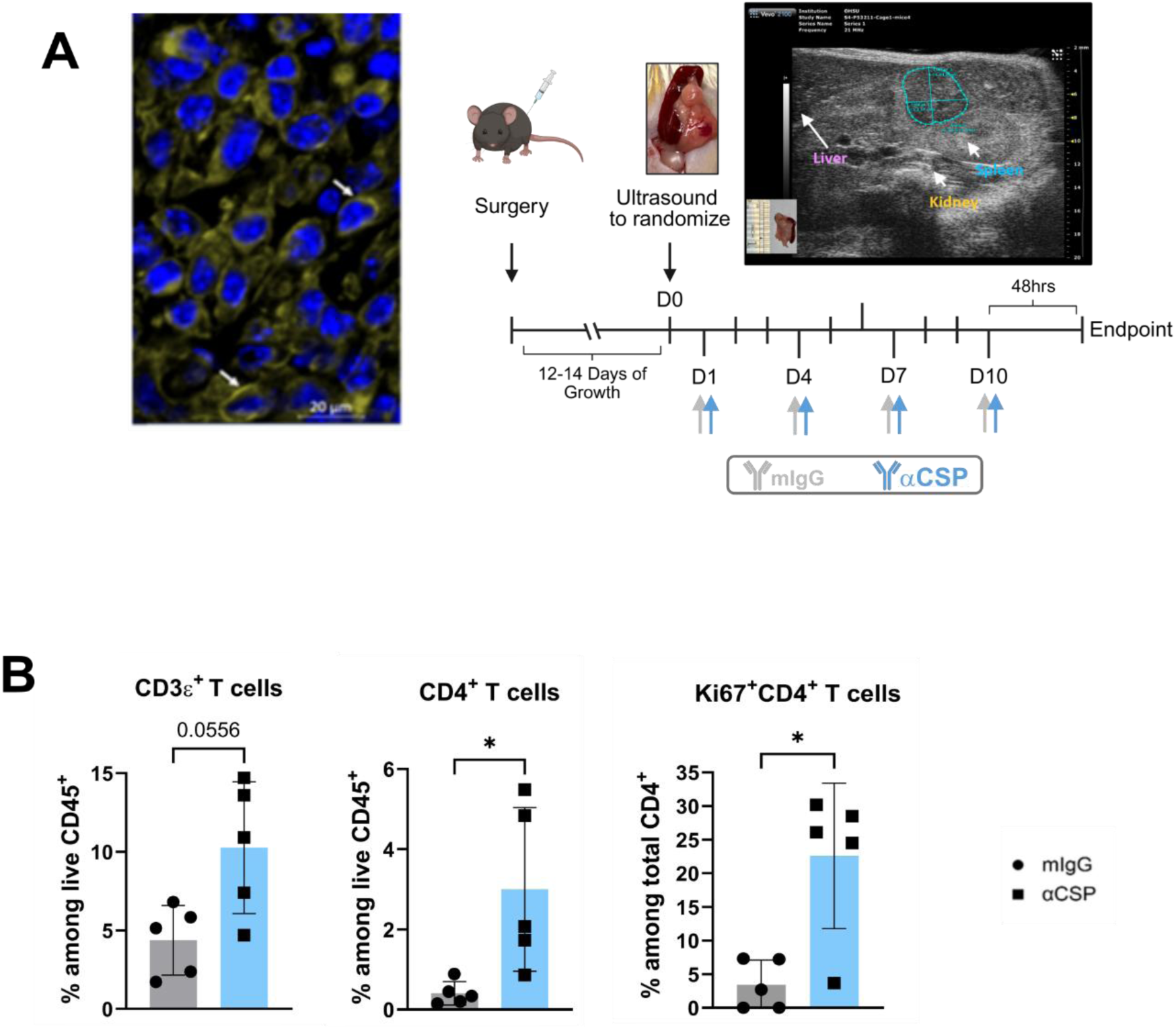
Orthotopic PDAC model shows increased cytotoxic T cell infiltration post αCSP treatment. **(A**) Treatment schema and ultrasound image showing tumor dimensions at beginning of treatment. **(B)** CD3^+^, CD4^+^ and Ki67^+^ CD4^+^ cells after treatment of either mIgG or αCSP. Wilcoxon rank-sum test. *p<0.05.

**Supplemental Table 2:** Antibodies used for multi-color flow cytometry.

| Fluorochrome | Marker | Dilution | Company | Cat# | RRID |
| --- | --- | --- | --- | --- | --- |
| BUV395 | CD45 | 1000 | BD Bioscience | 564279 | RRID:AB_2651134 |
| BUV661 | CD19 | 200 | BD Bioscience | 612971 | RRID:AB_2870243 |
| BUV737 | CD3 | 400 | BD Bioscience | 612803 | RRID:AB_2870130 |
| BUV805 | CD8 | 400 | BD Bioscience | 612898 | RRID:AB_2870186 |
| Super Bright 436 | Siglec-F | 200 | eBioscience | 62-1702-82 | RRID:AB_2664259 |
| eFluor 450 | MHCII | 1000 | Thermofisher Scientific | 48-5321-82 | RRID:AB_1272204 |
| BV480 | *Ki67* | 200 | BD Bioscience | 566109 | RRID:AB_2739511 |
| eFluor 506 | *FoxP3* | 100 | Thermofisher Scientific | 69-5773-82 | RRID:AB_2637367 |
| BV570 | Ly6C | 400 | Biolegend | 128029 | RRID:AB_10896061 |
| BV605 | CD11c | 400 | Biolegend | 117334 | RRID:AB_2562415 |
| BV650 | NK1.1 | 50 | Biolegend | 108735 | RRID:AB_11147949 |
| BV750 | CD103 | 200 | BD Bioscience | 747478 | RRID:AB_2872154 |
| Alexa Fluor 532 | CD11b | 400 | Thermofisher Scientific | 58-0112-82 | RRID:AB_2811905 |
| PE-Cy5.5 | *GrazB* | 100 | Invitrogen | 35889882 | RRID:AB_2848329 |
| Spark YG593 | CD44 | 200 | Biolegend | 103078 | RRID:AB_2892266 |
| PE-CF594 | PD-1 | 200 | BD Bioscience | 562523 | RRID:AB_2737634 |
| APC | CD206 | 100 | Biolegend | 141708 | RRID:AB_10900231 |
| Spark NIR 685 | Ly6G | 100 | Biolegend | 127666 | RRID:AB_2876454 |
| Alexa Fluor 700 | CCR7 | 100 | Thermofisher Scientific | 56-1971-82 | RRID:AB_657687 |
| APC-Fire 750 | F4/80 | 100 | Biolegend | 123152 | RRID:AB_2616725 |
| APC/Fire™ 810 | CD4 | 200 | Biolegend | 100480 | RRID:AB_2860583 |

## Declarations

### Ethics approval and consent to participate

All animal studies were approved by the University of Virginia and OHSU’s Institutional Animal Care and Use Committees

### Consent for publication

Not applicable

### Availability of data and materials

The datasets used and analyzed in this study are publicly available at the following locations:

Bulk RNA-sequencing data is available at the Genomic Database Commons website (https://portal.gdc.cancer.gov/). Single-cell RNA-sequencing data is available at https://zenodo.org/records/14199536. All code generated in this project to analyze data is available at https://github.com/CodyWolf1/Plectin-represses-cytotoxic-T-cell-activity-in-pancreatic-cancer.git.

### Authors’ contributions

**C.L. Wolf:** conceptualization, methodology, data curation, investigation, visualization, writing - original draft, writing – review and editing. **R.K.Ruiz**: conceptualization, methodology, data curation, investigation, visualization, writing – review and editing. **S.Khou**: methodology, investigation, data curation, visualization, writing – review and editing. **R.Cornelison**: methodology, writing – review and editing. **E.B.Stelow**: data curation. **K.M.Kowalewski**: methodology, visualization. **M.J.Lazzara**: methodology, visualization. **A.Poissonnier**: data curation, investigation. **L.M.Coussens**: methodology, supervision, funding acquisition, writing – review and editing. **K.A.Kelly**: conceptualization, methodology, supervision, funding acquisition, writing – original draft, writing – review and editing.

## Acknowledgements

Images detailing treatment schemas were created with BioRender.com

## References

1. R. L. Siegel, T. B. Kratzer, N. S. Wagle, H. Sung, A. Jemal, Cancer statistics, 2026. CA A Cancer J Clinicians 76, e70043 (2026).

2. L. Rahib, B. D. Smith, R. Aizenberg, A. B. Rosenzweig, J. M. Fleshman, L. M. Matrisian, Projecting Cancer Incidence and Deaths to 2030: The Unexpected Burden of Thyroid, Liver, and Pancreas Cancers in the United States. Cancer Research 74, 2913–2921 (2014).

3. C. Deiana, M. Agostini, G. Brandi, E. Giovannetti, The trend toward more target therapy in pancreatic ductal adenocarcinoma. Expert Review of Anticancer Therapy 24, 525–565 (2024).

4. Z. I. Hu, E. M. O’Reilly, Therapeutic developments in pancreatic cancer. Nat Rev Gastroenterol Hepatol 21, 7–24 (2024).

5. C. Peng, P. E. Oberstein, Emerging Therapeutic Approaches to Pancreatic Adenocarcinoma: Advances and Future Directions. Curr. Treat. Options in Oncol. (2025), doi:10.1007/s11864-025-01352-2.

6. T. F. Stoop, A. A. Javed, A. Oba, B. G. Koerkamp, T. Seufferlein, J. W. Wilmink, M. G. Besselink, Pancreatic cancer. The Lancet 405, 1182–1202 (2025).

7. H. Mi, S. Sivagnanam, C. B. Betts, S. M. Liudahl, E. M. Jaffee, L. M. Coussens, A. S. Popel, Quantitative Spatial Profiling of Immune Populations in Pancreatic Ductal Adenocarcinoma Reveals Tumor Microenvironment Heterogeneity and Prognostic Biomarkers. Cancer Research 82, 4359–4372 (2022).

8. J. Kleeff, M. Korc, M. Apte, C. La Vecchia, C. D. Johnson, A. V. Biankin, R. E. Neale, M. Tempero, D. A. Tuveson, R. H. Hruban, J. P. Neoptolemos, Pancreatic cancer. Nat Rev Dis Primers 2, 16022 (2016).

9. K. A. Kelly, N. Bardeesy, R. Anbazhagan, S. Gurumurthy, J. Berger, H. Alencar, R. A. DePinho, U. Mahmood, R. Weissleder, S. Gambhir, Ed. Targeted Nanoparticles for Imaging Incipient Pancreatic Ductal Adenocarcinoma. PLoS Med 5, e85 (2008).

10. G. Wiche, Plectin-Mediated Intermediate Filament Functions: Why Isoforms Matter. Cells 10, 2154 (2021).

11. M. J. Castañón, G. Walko, L. Winter, G. Wiche, Plectin–intermediate filament partnership in skin, skeletal muscle, and peripheral nerve. Histochem Cell Biol 140, 33–53 (2013).

12. A. C. Raymond, B. Gao, L. Girard, J. D. Minna, D. Gomika Udugamasooriya, Unbiased peptoid combinatorial cell screen identifies plectin protein as a potential biomarker for lung cancer stem cells. Sci Rep 9, 14954 (2019).

13. M. Buckup, M. A. Rice, E.-C. Hsu, F. Garcia-Marques, S. Liu, M. Aslan, A. Bermudez, J. Huang, S. J. Pitteri, T. Stoyanova, Plectin is a regulator of prostate cancer growth and metastasis. Oncogene 40, 663–676 (2021).

14. H. Gundesli, M. Kori, K. Y. Arga, The Versatility of Plectin in Cancer: A Pan-Cancer Analysis on Potential Diagnostic and Prognostic Impacts of Plectin Isoforms. OMICS: A Journal of Integrative Biology 27, 281–296 (2023).

15. K. Gao, Z. Gao, M. Xia, H. Li, J. Di, Role of plectin and its interacting molecules in cancer. Med Oncol 40, 280 (2023).

16. Z. Wang, W. Wang, Q. Luo, G. Song, Plectin: Dual Participation in Tumor Progression. Biomolecules 14, 1050 (2024).

17. G. Zhou, Y. Zhang, Z. Cai, H. Yao, M. Liu, C. Jiang, Z. Cheng, A biomimetic dual-targeting nanomedicine for pancreatic cancer therapy. J. Mater. Chem. B 13, 3716–3729 (2025).

18. Z. Outla, G. Oyman-Eyrilmez, K. Korelova, M. Prechova, L. Frick, L. Sarnova, P. Bisht, P. Novotna, J. Kosla, P. Bortel, Y. Borutzki, A. Bileck, C. Gerner, M. Rahbari, N. Rahbari, E. Birgin, B. Kvasnicova, A. Galisova, K. Sulkova, A. Bauer, N. Jobe, O. Tolde, E. Sticova, D. Rösel, T. O’Connor, M. Otahal, D. Jirak, M. Heikenwälder, G. Wiche, S. M. Meier-Menches, M. Gregor, Plectin-mediated cytoskeletal crosstalk as a target for inhibition of hepatocellular carcinoma growth and metastasis. eLife 13, RP102205 (2025).

19. S. M. Perez, J. Dimastromatteo, C. N. Landen, K. A. Kelly, A Novel Monoclonal Antibody Targeting Cancer-Specific Plectin Has Potent Antitumor Activity in Ovarian Cancer. Cells 10, 2218 (2021).

20. S. M. Perez, L. T. Brinton, K. A. Kelly, Plectin in Cancer: From Biomarker to Therapeutic Target. Cells 10, 2246 (2021).

21. D. Bausch, S. Thomas, M. Mino-Kenudson, C. C. Fernández-del, T. W. Bauer, M. Williams, A. L. Warshaw, S. P. Thayer, K. A. Kelly, Plectin-1 as a novel biomarker for pancreatic cancer. Clin Cancer Res 17, 302–309 (2011).

22. S. M. Perez, J. Dimastromatteo, C. N. Landen, K. A. Kelly, A Novel Monoclonal Antibody Targeting Cancer-Specific Plectin Has Potent Antitumor Activity in Ovarian Cancer. Cells 10, 2218 (2021).

23. O. G. Rikardsen, S. N. Magnussen, G. Svineng, E. Hadler-Olsen, L. Uhlin-Hansen, S. E. Steigen, Plectin as a prognostic marker in non-metastatic oral squamous cell carcinoma. BMC Oral Health 15, 98 (2015).

24. K. Katada, T. Tomonaga, M. Satoh, K. Matsushita, Y. Tonoike, Y. Kodera, T. Hanazawa, F. Nomura, Y. Okamoto, Plectin promotes migration and invasion of cancer cells and is a novel prognostic marker for head and neck squamous cell carcinoma. J Proteomics 75, 1803–1815 (2012).

25. S. J. Shin, J. A. Smith, G. A. Rezniczek, S. Pan, R. Chen, T. A. Brentnall, G. Wiche, K. A. Kelly, Unexpected gain of function for the scaffolding protein plectin due to mislocalization in pancreatic cancer. Proc Natl Acad Sci U S A 110, 19414–19419 (2013).

26. T. C. Burch, M. T. Watson, J. O. Nyalwidhe, Variable Metastatic Potentials Correlate with Differential Plectin and Vimentin Expression in Syngeneic Androgen Independent Prostate Cancer Cells. PLOS ONE 8, e65005 (2013).

27. C.-C. Cheng, W.-T. Chao, C.-C. Liao, Y.-H. Tseng, Y.-C. C. Lai, Y.-S. Lai, Y.-H. Hsu, Y.- H. Liu, Plectin deficiency in liver cancer cells promotes cell migration and sensitivity to sorafenib treatment. Cell Adh Migr 12, 19–27 (2018).

28. M. Žugec, B. Furlani, M. J. Castañon, B. Rituper, I. Fischer, G. Broggi, R. Caltabiano, G. M. V. Barbagallo, M. Di Rosa, D. Tibullo, R. Parenti, N. Vicario, S. Simčič, V. M. Pozo Devoto, G. B. Stokin, G. Wiche, J. Jorgačevski, R. Zorec, M. Potokar, Plectin plays a role in the migration and volume regulation of astrocytes: a potential biomarker of glioblastoma. Journal of Biomedical Science 31, 14 (2024).

29. R. L. Grossman, A. P. Heath, V. Ferretti, H. E. Varmus, D. R. Lowy, W. A. Kibbe, L. M. Staudt, Toward a Shared Vision for Cancer Genomic Data. N Engl J Med 375, 1109–1112 (2016).

30. R Core Team (2025), R: A Language and Environment for Statistical Computing_. R Foundation for Statistical Computing, Vienna, Austria. <https://www.R-project.org/>.

31. M. I. Love, W. Huber, S. Anders, Moderated estimation of fold change and dispersion for RNA-seq data with DESeq2. Genome Biol 15, 550 (2014).

32. A. Zhu, J. G. Ibrahim, M. I. Love, Heavy-tailed prior distributions for sequence count data: removing the noise and preserving large differences. Bioinformatics 35, 2084–2092 (2019).

33. R. Tibshirani, Estimating Transformations for Regression Via Additivity and Variance Stabilization. Journal of the American Statistical Association 83, 394–405 (1988).

34. W. Huber, A. von Heydebreck, H. Sueltmann, A. Poustka, M. Vingron, Parameter estimation for the calibration and variance stabilization of microarray data. Stat Appl Genet Mol Biol 2, Article3 (2003).

35. S. Anders, W. Huber, Differential expression analysis for sequence count data. Genome Biol 11, R106 (2010).

36. T. Wu, E. Hu, S. Xu, M. Chen, P. Guo, Z. Dai, T. Feng, L. Zhou, W. Tang, L. Zhan, X. Fu, S. Liu, X. Bo, G. Yu, clusterProfiler 4.0: A universal enrichment tool for interpreting omics data. Innovation (Camb) 2, 100141 (2021).

37. G. Yu, L.-G. Wang, Y. Han, Q.-Y. He, clusterProfiler: an R package for comparing biological themes among gene clusters. OMICS 16, 284–287 (2012).

38. A. Subramanian, P. Tamayo, V. K. Mootha, S. Mukherjee, B. L. Ebert, M. A. Gillette, A. Paulovich, S. L. Pomeroy, T. R. Golub, E. S. Lander, J. P. Mesirov, Gene set enrichment analysis: a knowledge-based approach for interpreting genome-wide expression profiles. Proc Natl Acad Sci U S A 102, 15545–15550 (2005).

39. A. Liberzon, A. Subramanian, R. Pinchback, H. Thorvaldsdóttir, P. Tamayo, J. P. Mesirov, Molecular signatures database (MSigDB) 3.0. Bioinformatics 27, 1739–1740 (2011).

40. J. Godec, Y. Tan, A. Liberzon, P. Tamayo, S. Bhattacharya, A. J. Butte, J. P. Mesirov, W. N. Haining, Compendium of Immune Signatures Identifies Conserved and Species-Specific Biology in Response to Inflammation. Immunity 44, 194–206 (2016).

41. M. Ashburner, C. A. Ball, J. A. Blake, D. Botstein, H. Butler, J. M. Cherry, A. P. Davis, K. Dolinski, S. S. Dwight, J. T. Eppig, M. A. Harris, D. P. Hill, L. Issel-Tarver, A. Kasarskis, S. Lewis, J. C. Matese, J. E. Richardson, M. Ringwald, G. M. Rubin, G. Sherlock, Gene Ontology: tool for the unification of biology. Nat Genet 25, 25–29 (2000).

42. M. Kanehisa, M. Furumichi, Y. Sato, M. Kawashima, M. Ishiguro-Watanabe, KEGG for taxonomy-based analysis of pathways and genomes. (available at 10.1093/nar/gkac963).

43. M. Milacic, D. Beavers, P. Conley, C. Gong, M. Gillespie, J. Griss, R. Haw, B. Jassal, L. Matthews, B. May, R. Petryszak, E. Ragueneau, K. Rothfels, C. Sevilla, V. Shamovsky, R. Stephan, K. Tiwari, T. Varusai, J. Weiser, A. Wright, G. Wu, L. Stein, H. Hermjakob, P. D’Eustachio, The Reactome Pathway Knowledgebase 2024. (available at 10.1093/nar/gkad1025).

44. E. Ulgen, O. Ozisik, O. U. Sezerman, pathfindR: An R Package for Comprehensive Identification of Enriched Pathways in Omics Data Through Active Subnetworks. Front Genet 10, 858 (2019).

45. D. W. Huang, B. T. Sherman, Q. Tan, J. R. Collins, W. G. Alvord, J. Roayaei, R. Stephens, M. W. Baseler, H. C. Lane, R. A. Lempicki, The DAVID Gene Functional Classification Tool: a novel biological module-centric algorithm to functionally analyze large gene lists. Genome Biol 8, R183 (2007).

46. Aragaki K, Yoshihara K, Vegesna R, Kim H, Verhaak R, tidyestimate: A Tidy Implementation of “ESTIMATE” (available at https://github.com/kaiaragaki/tidyestimate.).

47. A. M. Newman, C. B. Steen, C. L. Liu, A. J. Gentles, A. A. Chaudhuri, F. Scherer, M. S. Khodadoust, M. S. Esfahani, B. A. Luca, D. Steiner, M. Diehn, A. A. Alizadeh, Determining cell type abundance and expression from bulk tissues with digital cytometry. Nat Biotechnol 37, 773–782 (2019).

48. H. Wickham, ggplot2 (Springer International Publishing, Cham, 2016; http://link.springer.com/10.1007/978-3-319-24277-4).

49. I. M. Loveless, S. B. Kemp, K. M. Hartway, J. T. Mitchell, Y. Wu, S. D. Zwernik, D. J. Salas-Escabillas, S. Brender, M. George, Y. Makinwa, T. Stockdale, K. Gartrelle, R. G. Reddy, D. W. Long, A. Wombwell, J. M. Clark, A. M. Levin, D. Kwon, L. Huang, R. Francescone, D. B. Vendramini-Costa, B. Z. Stanger, A. Alessio, A. M. Waters, Y. Cui, E. J. Fertig, L. T. Kagohara, B. Theisen, H. C. Crawford, N. G. Steele, Human Pancreatic Cancer Single-Cell Atlas Reveals Association of CXCL10+ Fibroblasts and Basal Subtype Tumor Cells. Clin Cancer Res 31, 756–772 (2025).

50. Y. Hao, T. Stuart, M. H. Kowalski, S. Choudhary, P. Hoffman, A. Hartman, A. Srivastava, G. Molla, S. Madad, C. Fernandez-Granda, R. Satija, Dictionary learning for integrative, multimodal and scalable single-cell analysis. Nat Biotechnol 42, 293–304 (2024).

51. T. Stuart, A. Butler, P. Hoffman, C. Hafemeister, E. Papalexi, W. M. Mauck, Y. Hao, M. Stoeckius, P. Smibert, R. Satija, Comprehensive integration of single-cell data. Cell 177, 1888–1902.e21 (2019).

52. L. McInnes, J. Healy, J. Melville, UMAP: Uniform Manifold Approximation and Projection for Dimension Reduction (2020), doi:10.48550/arXiv.1802.03426.

53. J. W. Squair, M. Gautier, C. Kathe, M. A. Anderson, N. D. James, T. H. Hutson, R. Hudelle, T. Qaiser, K. J. E. Matson, Q. Barraud, A. J. Levine, G. La Manno, M. A. Skinnider, G. Courtine, Confronting false discoveries in single-cell differential expression. Nat Commun 12, 5692 (2021).

54. H. L. Crowell, C. Soneson, P.-L. Germain, D. Calini, L. Collin, C. Raposo, D. Malhotra, M. D. Robinson, muscat detects subpopulation-specific state transitions from multi-sample multi-condition single-cell transcriptomics data. Nat Commun 11, 6077 (2020).

55. J.-C. Beltra, S. Manne, M. S. Abdel-Hakeem, M. Kurachi, J. R. Giles, Z. Chen, V. Casella, S. F. Ngiow, O. Khan, Y. J. Huang, P. Yan, K. Nzingha, W. Xu, R. K. Amaravadi, X. Xu, G. C. Karakousis, T. C. Mitchell, L. M. Schuchter, A. C. Huang, E. J. Wherry, Developmental Relationships of Four Exhausted CD8+ T Cell Subsets Reveals Underlying Transcriptional and Epigenetic Landscape Control Mechanisms. Immunity 52, 825–841.e8 (2020).

56. B. C. Miller, D. R. Sen, R. Al Abosy, K. Bi, Y. V. Virkud, M. W. LaFleur, K. B. Yates, A. Lako, K. Felt, G. S. Naik, M. Manos, E. Gjini, J. R. Kuchroo, J. J. Ishizuka, J. L. Collier, G. K. Griffin, S. Maleri, D. E. Comstock, S. A. Weiss, F. D. Brown, A. Panda, M. D. Zimmer, R. T. Manguso, F. S. Hodi, S. J. Rodig, A. H. Sharpe, W. N. Haining, Subsets of exhausted CD8+ T cells differentially mediate tumor control and respond to checkpoint blockade. Nat Immunol 20, 326–336 (2019).

57. S. Cheng, Z. Li, R. Gao, B. Xing, Y. Gao, Y. Yang, S. Qin, L. Zhang, H. Ouyang, P. Du, L. Jiang, B. Zhang, Y. Yang, X. Wang, X. Ren, J.-X. Bei, X. Hu, Z. Bu, J. Ji, Z. Zhang, A pan-cancer single-cell transcriptional atlas of tumor infiltrating myeloid cells. Cell 184, 792–809.e23 (2021).

58. I. Tirosh, B. Izar, S. M. Prakadan, M. H. Wadsworth, D. Treacy, J. J. Trombetta, A. Rotem, C. Rodman, C. Lian, G. Murphy, M. Fallahi-Sichani, K. Dutton-Regester, J.-R. Lin, O. Cohen, P. Shah, D. Lu, A. S. Genshaft, T. K. Hughes, C. G. K. Ziegler, S. W. Kazer, A. Gaillard, K. E. Kolb, A.-C. Villani, C. M. Johannessen, A. Y. Andreev, E. M. Van Allen, M. Bertagnolli, P. K. Sorger, R. J. Sullivan, K. T. Flaherty, D. T. Frederick, J. Jané-Valbuena, C. H. Yoon, O. Rozenblatt-Rosen, A. K. Shalek, A. Regev, L. A. Garraway, Dissecting the multicellular ecosystem of metastatic melanoma by single-cell RNA-seq. Science 352, 189–196 (2016).

59. S. R. Hingorani, L. Wang, A. S. Multani, C. Combs, T. B. Deramaudt, R. H. Hruban, A. K. Rustgi, S. Chang, D. A. Tuveson, *Trp53R172H* and *KrasG12D* cooperate to promote chromosomal instability and widely metastatic pancreatic ductal adenocarcinoma in mice. Cancer Cell 7, 469–483 (2005).

60. K. Kelly, R. Burton, A. Horwitz, Cancer Specific Plectin-1 Specific Antibodies and Methods of Use Thereof (2023).

61. G. L. Beatty, E. G. Chiorean, M. P. Fishman, B. Saboury, U. R. Teitelbaum, W. Sun, R. D. Huhn, W. Song, D. Li, L. L. Sharp, D. A. Torigian, P. J. O’Dwyer, R. H. Vonderheide, CD40 Agonists Alter Tumor Stroma and Show Efficacy Against Pancreatic Carcinoma in Mice and Humans. Science 331, 1612–1616 (2011).

62. A. Weiss, D. R. Littman, Signal transduction by lymphocyte antigen receptors. Cell 76, 263–274 (1994).

63. J. L. Benci, B. Xu, Y. Qiu, T. J. Wu, H. Dada, C. Twyman-Saint Victor, L. Cucolo, D. S. M. Lee, K. E. Pauken, A. C. Huang, T. C. Gangadhar, R. K. Amaravadi, L. M. Schuchter, M. D. Feldman, H. Ishwaran, R. H. Vonderheide, A. Maity, E. J. Wherry, A. J. Minn, Tumor Interferon Signaling Regulates a Multigenic Resistance Program to Immune Checkpoint Blockade. Cell 167, 1540–1554.e12 (2016).

64. J. L. Benci, L. R. Johnson, R. Choa, Y. Xu, J. Qiu, Z. Zhou, B. Xu, D. Ye, K. L. Nathanson, C. H. June, E. J. Wherry, N. R. Zhang, H. Ishwaran, M. D. Hellmann, J. D. Wolchok, T. Kambayashi, A. J. Minn, Opposing Functions of Interferon Coordinate Adaptive and Innate Immune Responses to Cancer Immune Checkpoint Blockade. Cell 178, 933–948.e14 (2019).

65. M. R. Goulart, K. Stasinos, R. E. A. Fincham, F. R. Delvecchio, H. M. Kocher, T cells in pancreatic cancer stroma. World Journal of Gastroenterology 27, 7956 (2021).

66. D. H. Munn, A. L. Mellor, IDO in the Tumor Microenvironment: Inflammation, Counter-Regulation, and Tolerance. Trends Immunol 37, 193–207 (2016).

67. L. Zitvogel, L. Galluzzi, O. Kepp, M. J. Smyth, G. Kroemer, Type I interferons in anticancer immunity. Nat Rev Immunol 15, 405–414 (2015).

68. M. R. Goulart, K. Stasinos, R. E. A. Fincham, F. R. Delvecchio, H. M. Kocher, T cells in pancreatic cancer stroma. World J Gastroenterol 27, 7956–7968 (2021).

69. J. Ge, J. Ge, G. Tang, D. Xiong, D. Zhu, X. Ding, X. Zhou, M. Sang, Machine learning-based identification of biomarkers and drugs in immunologically cold and hot pancreatic adenocarcinomas. J Transl Med 22, 775 (2024).

